# Soluble DLK1 secreted by telomere-shortening-induced senescent microglia impairs oligodendrocyte functions and alters neuronal activity

**DOI:** 10.64898/2026.01.14.699608

**Authors:** Bangyan Liu, Matthew Mahoney, Yilin Feng, Maria Telpoukhovskaia, Alice Maria Giani, Eileen Ruth Torres, Lihong Zhan, Pearly Ye, Jingjie Zhu, Nessa R Foxe, Daphne Zhu, Xinran Tong, Deepak Srivastava, Christine V. Theodoris, Shiaoching Gong, Mingrui Zhao, Li Fan, Li Gan

## Abstract

Aging is the major risk factor for neurodegenerative disease, yet the mechanisms linking physiological aging to brain dysfunction remain unclear. Because telomere erosion is a hallmark of aging, we examined its impact on glial and neuronal physiology. Telomere shortened mice showed lipofuscinosis, hypomyelination, microglial atrophy, and cognitive deficits. Single nuclei RNA-seq revealed accelerated glial aging, elevated microglial senescence pathways, and impaired oligodendrocyte functions. Inducing senescence in human iPSC derived microglia with shortened telomeres identified soluble DLK1 as a novel senescence associated ligand. sDLK1 was increased in the cerebrospinal fluid of telomere shortened and naturally aged mice, and this increase was eliminated by microglial depletion. AAV delivery of sDLK1 in vivo caused hypomyelination and blocked oligodendrocyte lineage progression, demonstrating the detrimental nature of excessive sDLK1. In human iPSC systems, sDLK1 impaired oligodendrocyte maturation and altered calcium signaling in excitatory neurons. These findings identify microglial senescence as a core consequence of telomere shortening and reveal sDLK1 as a microglia-derived senescence ligand that drives oligodendrocyte and neuronal dysfunction in aging.

## Introduction

Aging is the most critical risk factor for many neurodegenerative disorders, including Alzheimer’s disease (AD), the most common cause of dementia, which contributes to up to 80% of all dementia cases^1^. Rather than presenting with a single neurodegenerative pathology, a large proportion of the elderly dementia patients exhibit mixed brain pathologies, including AD and other degenerative and vascular brain pathologies simultaneously^2,3^. The comprehensive disruption of brain functions and structural integrity can be attributed to pathological brain aging, which is characterized by more rapid deterioration due to genetic predisposition and environmental factors^4–6^. Understanding the mechanisms underlying pathological brain aging is critical for unveiling the pathogenesis of neurodegenerative disorders and other age-dependent brain conditions.

Telomeres are repeated nucleotide sequences at the ends of chromosomes that prevent nucleolytic degradation and chromosome end-to-end fusion. The telomere is a key indicator of an organism’s biological age^7^. Human telomeres range from 2 to 30 kb and gradually lose their length due to the end replication problem during aging^8,9^. Telomere sequence and function are conserved between human and mouse, but mouse telomeres are 5–10 times longer than human telomeres, and mouse lifespan is 30 times shorter, making it unfeasible to study the mechanisms affected by critically shortened telomeres in naturally aged mice^10^. To overcome this, TERC knock-out has been used to study the effect of telomere shortening in mice. TERC is the RNA component of telomerase, an RNA-protein complex that elongates telomere length in cells with infinite proliferative potential and serves as the replication template for telomere sequence^11^. Knockout of TERC hampers the telomerase activity without affecting other enzymatic functions of TERT, the protein component of telomerase, and causes telomere shortening to be inherited through generations by disabling telomere elongations in the mouse germline. Previous studies have showcased that three consecutive generations of TERC knock-out induce aging-like phenotypes^12^.

Critical shortening of telomeres by the end replication problem induces cell-cycle arrest and causes the cell to enter replicative senescence^13^. Glial cells in the brain (e.g., microglia, astrocytes, and oligodendrocytes) retain proliferative capabilities after development and become more proliferative in response to damage to the central nervous system (CNS) and other stressors. Thus, glial cells are under heavy replicative stress, and telomere shortening is detected in the white matter, whereas telomere length in the grey matter remains relatively unchanged^14,15^.

Glial senescence has been suggested to transform the brain from normal aging to pathological aging and to drive the buildup and spread of AD pathologies^16^. Microglia are the resident macrophages in the CNS responsible for immune surveillance and innate immune responses to damage and pathogenic species^17^. Microglia are susceptible to more replicative stress associated with the reactivation of the proliferative program caused by the responses to neurodegenerative pathologies, including tauopathy and Aβ accumulation^18,19^. Telomere shortening and replicative senescence may disrupt normal microglia functions under aging and neurodegenerative conditions^20–22^. However, how microglia senescence contributes to pathological aging and affects other brain cell types remains unknown.

To determine the effects of telomere shortening in the brain, we assessed pathological and functional outcomes while characterizing the cell-type-specific transcriptomic alterations. We expanded our study on telomere-shortened mouse microglia to human iPSC-derived microglia with shortened telomeres. Here, we report direct evidence that senescent microglia exert detrimental influences on other cell types through an altered secretion profile. Importantly, we identified delta-like non-canonical Notch ligand 1 (DLK1) as a new senescent microglia-derived ligand. DLK1 was originally identified as a member of the epidermal growth factor (EGF)-like family and a negative regulator of adipocyte differentiation (pre-adipocyte inhibitor factor 1, Pref1)^23^. Overexpression of soluble DLK1 (sDLK1) in the mouse brain caused defects in oligodendrocyte differentiation and myelinating activities. sDLK1 treatment of human iPSC-derived oligodendrocytes blocked normal maturation with expansion of OPCs, providing evidence that sDLK1 impairs oligodendrocytes directly. Higher levels of sDLK1 also induced abnormal Ca^2+^ activities in the mouse and human iPSC-derived neurons. This study profiles cellular signatures associated with shorter telomeres in the mouse brain and identifies replicative microglia senescence along with the associated ligands as a key contributor to pathological brain aging.

## Results

### Telomere shortening exacerbates brain lipofuscinosis

To understand the effects of telomere shortening on brain functions, we used the TERC knockout model. We bred mice with a homologous TERC deletion for three consecutive generations (G3 *Terc^-/-^*) and, at 8 months of age, conducted all subsequent analyses (**Fig. 1A**). *Terc^-/-^*mice have been shown to exhibit progressive telomere loss with increasing generations^24^. Using *in situ* hybridization, we measured the fluorescence intensities of a probe complementary to the telomere sequence in the mouse cortex, which directly correlate with telomere length (**Fig. 1B**)^25^. As expected, quantification of the fluorescence intensity showed that G3 *Terc^-/-^* mice exhibited significantly less fluorescence in all and Iba1-positive cells than age-matched wild-type (WT) mice (**Fig. 1C, D**) in both female (**Fig. 1E**) and male (**Fig. 1F**) mice.

**Fig. 1.**
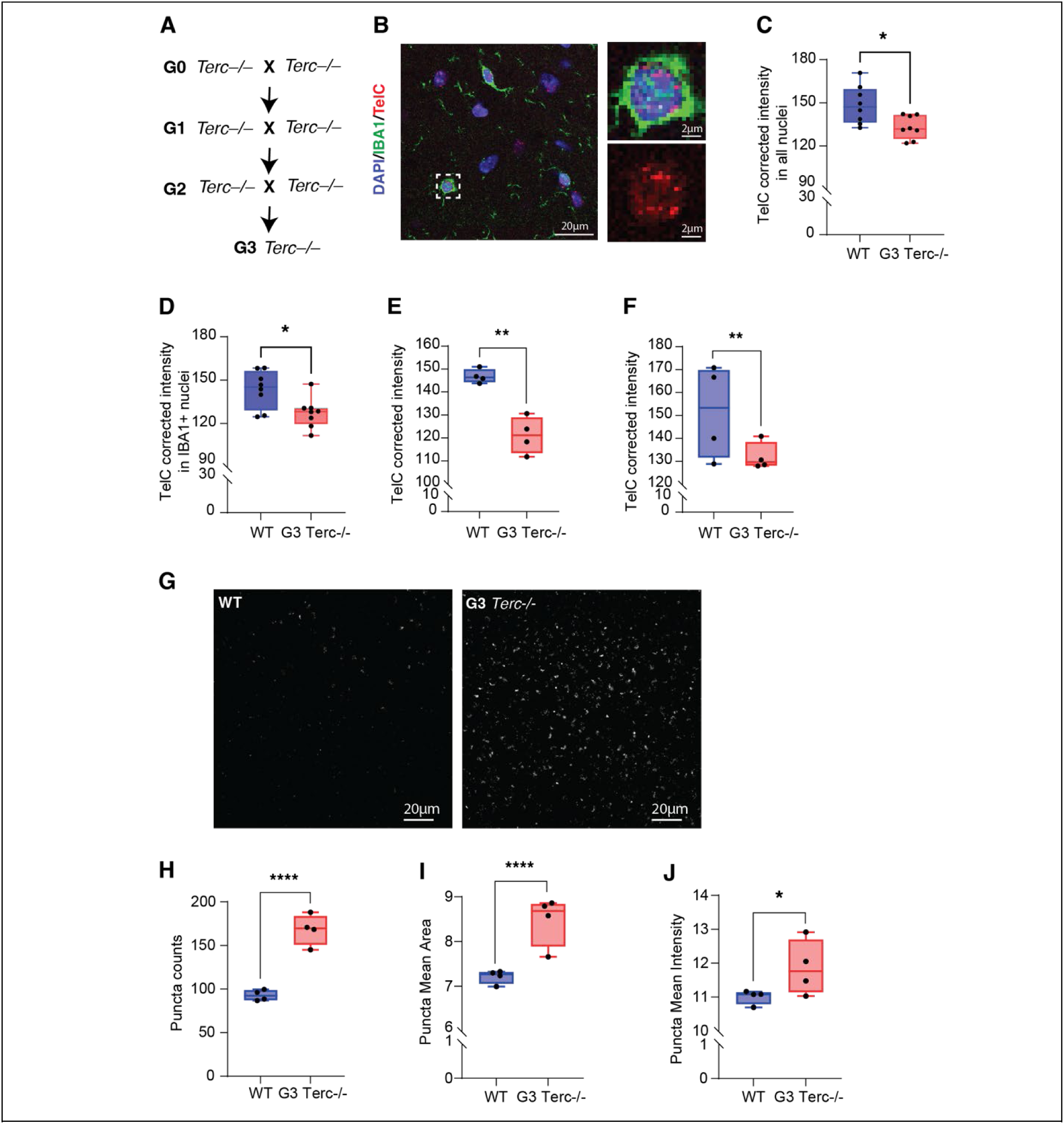
Telomere shortening exacerbates brain lipofuscinosis. **A.** Schematic showing the generation of G3 TERC-/- mice. **B.** Representative immunofluorescence images of telomere length measurements by *in situ* hybridization with TelC-Cy3 (red), DAPI (blue), and anti-IBA1 (green). Zoomed in images on the top right and TelC-Cy3-only image on the bottom right. 8 animals were tested per genotype; 3 sections were used for each animal, and 3 images were taken per section. ∼40 cells were captured in each image. **C.** Quantification of TelC-Cy3 fluorescence intensities in all cells (IBA1+ & IBA1-), showing reduced telomere length in the G3 Terc-/- brains. Results are presented as mean intensity measurements from eight animals. Data were analyzed by two-tailed unpaired *t*-test. \**p*=0.0133 **D.** Quantification of TelC-Cy3 fluorescence intensities in IBA1+ cells. Results are presented as mean intensity measurements from eight animals. Data were analyzed by two-tailed unpaired *t*-test. \**p*=0.016 **E–F.** Quantification of TelC-Cy3 fluorescence intensities in female (E) and male (F) mice. Four animals per genotype were analyzed, with 9 to 19 images per animal. Data are shown as box and whisker plots and analyzed by mixed model. \*\**p* = 0.0074 (female), \*\**p* = 0.007 (male). **G**. Representative autofluorescence images showing lipofuscin in the primary somatosensory cortex. **H-J.** Quantification of lipofuscin puncta counts (H), puncta mean area (I), puncta mean intensity (J). 4 animals were tested per genotype; 6-8 images were taken for each animal. Data were reported as box & whisker plot showing min to max and analyzed by mixed model. \*\*\*\**p*<0.0001, \**p*=0.0472.

Lipofuscinosis is an aging hallmark in the brain that can be visualized as autofluorescence^26^. Autofluorescence was strikingly increased throughout the brain, as detected in the primary somatosensory cortex (**Fig. 1G**). Quantification of the autofluorescence revealed a significant increase of lipofuscinosis in the G3 *Terc^-/-^*mice (**Fig. 1H–J**).

### Telomere shortening accelerates aging in glial cells

To investigate the mechanism by which telomere shortening affect brain functions, we performed snRNA-seq to examine hippocampal tissues from WT and G3 *Terc^-/-^* animals. We utilized a pre-established cold mechanical dissociation protocol to prepare the snRNA library and sequenced 83,223 nuclei^27^. Of these, 9,131 nuclei were filtered out, based on the gene counts, unique molecular identifier (UMI), and percent mitochondrial genes per nucleus (**Supp Fig. 1A-B**). We further removed potential multiplets identified by DoubletFinder and selected 72,330 nuclei for downstream analysis^28^. Graph-based clustering identified eight major cell types in the mouse hippocampus (**Fig. 2A**). Telomere shortening did not significantly affect the composition of the major cell types (**Supp Fig. 1C, D**). The cell types were annotated by known reference gene sets **(Supp Fig. 1E)**.

**Fig. 2.**
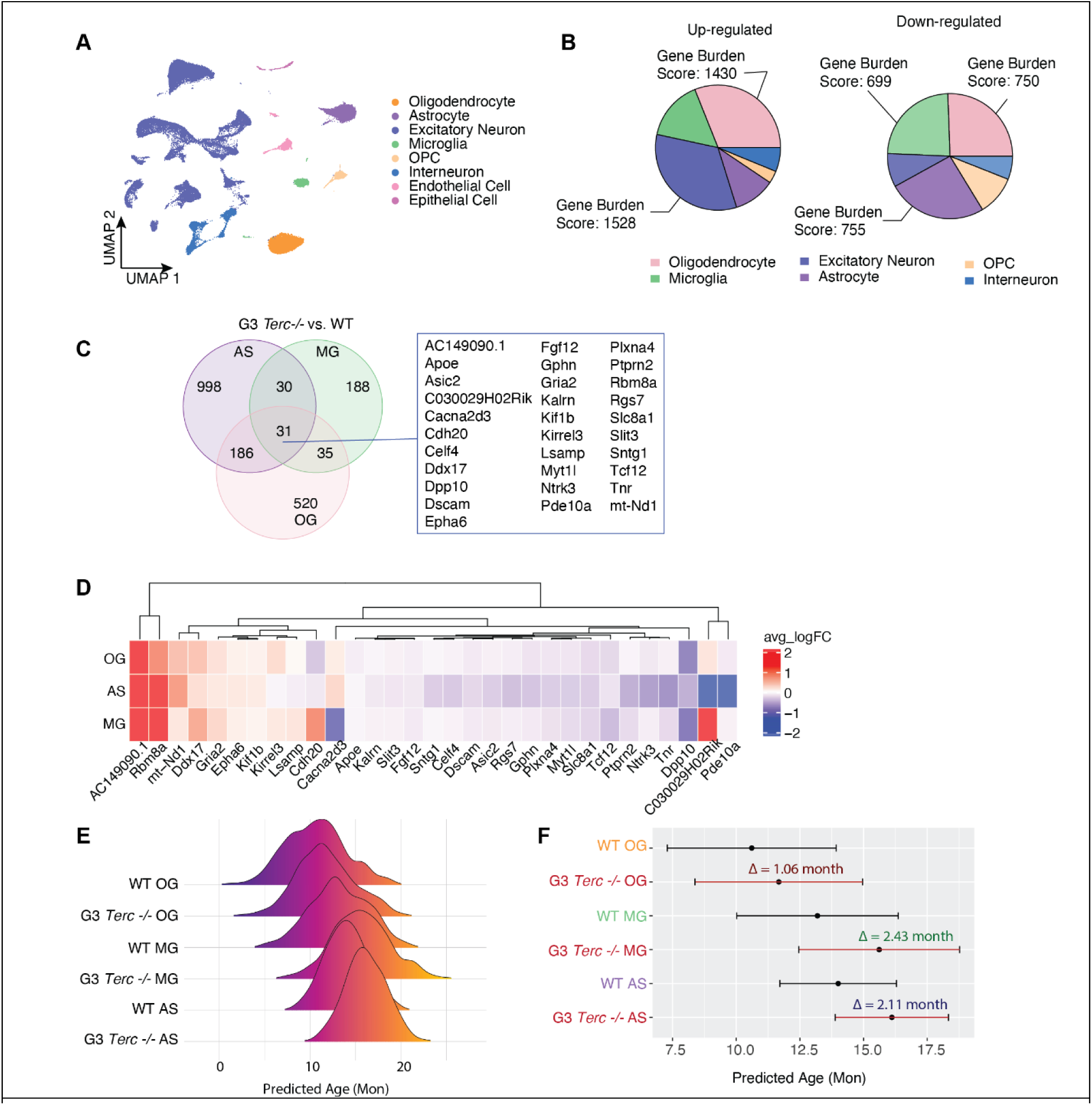
Telomere shortening accelerates aging in glial cells. **A.** Uniform Manifold Approximation and Projection (UMAP) plots showing eight major cell types identified in the mouse hippocampus. **B.** Distributions of the normalized numbers of genes up-regulated (top) and down-regulated (bottom) in different cell types of the G3 Terc^-/-^ mice. The gene burden score is defined as the number of differentially expressed genes per 1000 UMI detected in each cell type. **C.** Venn diagram showing numbers and overlaps of DEGs in oligodendrocyte, astrocyte, and microglia. DEGs common among the three cell types are listed in the rectangle on the right. **D.** Heatmap comparing the average log fold-change of the common DEGs in astrocytes, oligodendrocytes, and microglia in G3 Terc^-/-^ to WT mice. **E.** Ridge plot of the predicted chronological ages for oligodendrocyte, astrocyte, and microglia in G3 Terc^-/-^ and WT mice.

We focused on microglia, astrocytes, neurons, oligodendrocytes, and oligodendrocyte precursor cells for the downstream analysis. To identify the magnitude of telomere shortening effect on each cell type, we performed differential expression analysis for each cell type, comparing the transcriptomes of G3 *Terc^-/-^*and WT tissues. Telomere shortening led to significant alterations in all six analyzed cell types. The number of differentially expressed genes (DEGs) in each cell type was normalized, using the average number of UMIs detected for the cell type (**Fig. 2B**, **Supplementary Table 1**). Examination of the DEGs of microglia, astrocyte, and oligodendrocyte revealed that telomere shortening resulted in distinct transcriptomic changes in individual cell types (**Fig. 2C**). Nonetheless, we identified 31 DEGs shared among the three cell types, and the majority of the overlapped DEGs (26/31) were changed in the same direction by telomere shortening (**Fig. 2D**). Noticeably, AC149090.1 was significantly up-regulated in all three cell types. AC149090.1 is a mouse ortholog to human PISD, a gene that encodes for a phospholipid decarboxylase that catalyzes the conversion of phosphatidylserine to phosphatidylethanolamine in the inner mitochondrial membrane and is essential in lipid metabolism and autophagy^29^. AC149090.1 is a major marker of cell-type-specific aging in the neurogenic region of the subventricular zone^30^. Utilizing the aging clock built by the Brunet group^30^, we examined the effect of telomere shortening on predicted biological age in oligodendrocyte, microglia, and astrocytes (**Supp Fig. 1F**). Although the aging clocks are based on different gene contributors for different cell types (**Supp Fig. 1G**), we found that telomere shortening increased the predicted biological age in three glial cell types with microglia affected the most, accelerated by an estimated 2.43 months (**Fig. 2E, F**).

### Telomere shortening enhances senescence signaling in microglia and astrocytes and disrupts normal oligodendrocyte functions

We next analyzed transcriptomic changes induced by telomere shortening in microglia. The WT and G3 *Terc^-/-^* samples were subclustered into three distinct transcriptional states (**Fig. 3A**). A significant shift of microglia transcriptomic states was observed in microglia from WT and G3 *Terc ^-/-^* brains. Specifically, subcluster 3 (MG3) microglia in G3 *Terc^-/-^* animals were enriched (**Fig. 3B**). Applying the aging clock to MG3 showed that a subset was predicted to be markedly older than other microglia, linking telomere shortening induced transcriptomic changes to signatures of chronic brain aging (**Fig. 3C**).

**Fig. 3.**
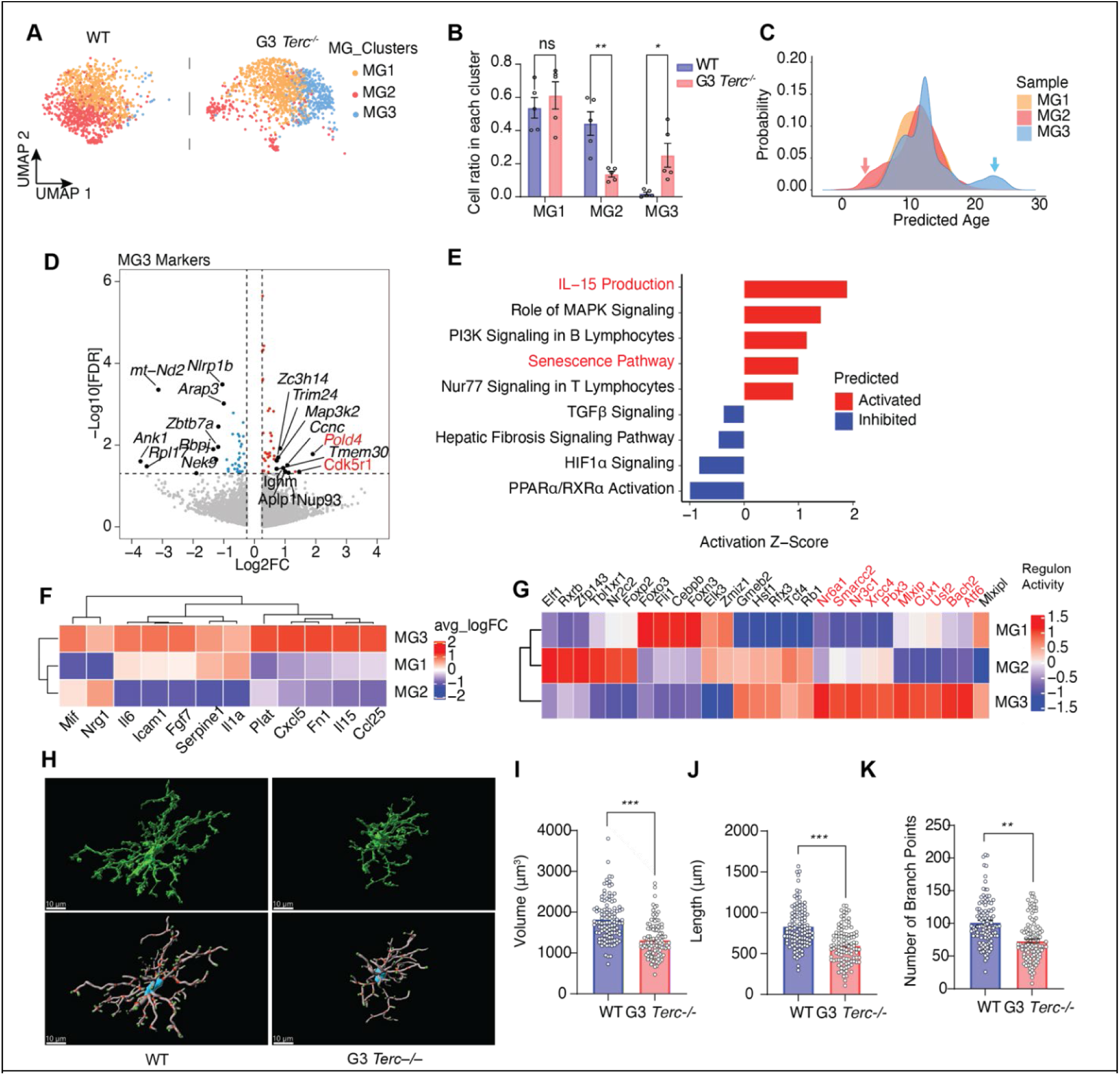
Telomere shortening induces microglial senescence pathways and atrophy. **A.** UMAP plot showing a shift from cluster 2 (red) to cluster 3 (blue) caused by G3 Terc^-/-^. **B.** Ratios of each microglia cluster. Data of each cluster were analyzed by Kruskal Willis test. \*\**p*=0.0013; \**p*=0.011 **C.** Density plot showing the predicted age of different microglia clusters. The red arrow points to a younger microglia population, and the blue arrow points at an aged microglia population. **D.** Volcano plot of marker genes of microglia cluster 3 (MG3). Red and blue dots represent genes with a log2FC > 0.1 or < -0.1, respectively. All other genes are colored gray. Top differential genes are labeled. **E.** Top canonical pathways identified by Ingenuity Pathway Analysis that are associated with DEGs in MG3. **F.** Heatmap showing the SASP genes upregulated in MG3. **G.** Heatmap showing the regulon activity predicted by SCENIC in individual microglia subclusters. **H.** (Top) Representative 3D Imaris surface renderings of microglia from 10-month-old WT and G3 Terc KO hippocampus. Scale bar represents 10 microns. (Bottom) Representative 3D Imaris filament tracings of microglia displaying soma, processes, branch points, and terminal points. Scale bar represents 10 microns. **I.** Quantification of microglia volume using Imaris 3D reconstruction. Each dot represents one microglia. n = 110 microglia for WT and n = 110 microglia for G3 Terc KO, where N = 5 mice/genotype and 3-4 hippocampal sections/mouse. Data are reported as mean ± SEM. \*\*\**p* = 0.0009. Data were analyzed by nested t-test/mixed model. **J.** Quantification of total process length/microglia using Imaris 3D reconstruction. Each dot represents one microglia. n = 110 microglia for WT and n = 110 microglia for G3 Terc KO, where N = 5 mice/genotype and 3-4 hippocampal sections/mouse. Data are reported as mean ± SEM. \*\*\**p* = 0.0007. Data were analyzed by nested t-test/mixed model. **K.** Quantification of number of branch points/microglia using Imaris 3D reconstruction. Each dot represents one microglia. n = 110 microglia for WT and n = 110 microglia for G3 Terc KO, where N = 5 mice/genotype and 3-4 hippocampal sections/mouse. Data are reported as mean ± SEM. \*\**p* = 0.002. Data were analyzed by nested t-test/mixed model.

Further analyses of top marker genes of MG3 revealed genes associated with replication stress survival and senescence, such as Pold4 and Cdk5r1 (**Fig. 3D**)^31,32^. Upregulated pathways identified in MG3 by Ingenuity Pathway Analysis (IPA) included the senescence pathway and production of IL-15, a known senescence-associated secretory phenotype (SASP) cytokine (**Fig. 3E**). We also examined expression changes of known SASP ligands and found 12 SASP ligands upregulated in MG3 (**Fig. 3F**). Single-cell regulatory network inference and clustering (SCENIC) predicted 10 upstream transcriptional factors specific to MG3 (**Fig. 3G**), including CUX1 and ATF6 involved in replicative senescence and morphological changes associated with senescence^33,34^.

Close examination revealed that microglia from G3 *Terc^-/-^*animals exhibited clear signs of morphological atrophy characteristic of aging (**Fig. 3H**). Specifically, G3 Terc^-/-^ microglia displayed significantly reduced cell body volume and shorter processes compared to the wild type (**Fig. 3I, J**), along with a marked decrease in the number of process branch points per cell (**Fig. 3K**). This retracted, less-ramified morphology is a well-established hallmark of aging microglia and reflects the dystrophic changes that accompany microglial senescence^35,36^. These structural alterations align with the senescent phenotype observed in telomere-shortened microglia.

Significant transcriptomic changes are also induced by telomere shortening in astrocytes, as two astrocyte subclusters, AS2 and AS6, are enriched in G3 *Terc^-/-^* animals (**Supp Fig. 2A, B**). Analysis of the DEGs distinguishing AS2 and AS6 from the homeostatic cluster AS1 showed that both subclusters were enriched for the SenMayo gene set, indicating that cellular senescence induced by telomere shortening is not limited to microglia (**Supp Fig. 2C**)^37^.

Loss of white matter volume is a well-documented phenomenon in the aging brain^38^. We examined the changes induced by telomere shortening in oligodendrocyte lineage cells by combining and re-clustering the oligodendrocyte precursor cells (OPCs) and oligodendrocytes in the UMAP (**Supp Fig. 2D**). We found that telomere shortening caused a striking shift from the OG2 to the OG1 subcluster (**Supp Fig. 2E**). Notably, genes associated with normal formation of myelin structures and neuroprotective effects, such as ANLN, SPOCK3, and APOD, were significantly reduced in OG1, the subcluster enriched in the G3 *Terc^-/-^* mice (**Supp Fig. 2F**)^39–41^. With OPC as a starting point, we performed pseudotime analyses to examine the maturation process from OPCs to oligodendrocytes. Telomere shortening shifted oligodendrocytes further along the pseudotime trajectory, indicating an abnormal maturation process that deviates from the typical lineage progression observed in wild-type cells (**Supp Fig. 2G**). Analyses of the expression dynamics of ANLN, SPOCK3, and APOD revealed a similar pattern, where the expression levels increased during the differentiation from the OPCs and reduced further away from the starting point, suggesting a connection between the disruption of normal myelin formation and increased pseudotime induced by telomere shortening (**Supp Fig. 2H**). To directly assess the impact of telomere shortening on oligodendrocyte functions, we conducted a western blot analysis and showed a marked reduction of myelin basic protein (MBP) in the cortex of G3 Terc–/– brains compared to wild-type controls (**Supp Fig. 2I**).

### Telomere shortening alters microglia-neuron interactions and induces cognitive deficits

We next examined the intercellular signaling between microglia and neuronal cells with CellChat. Among the 11 hippocampal neuronal subtypes in our snRNA dataset, we identified 268 total pathways from microglia to neurons that are common between the three microglial subclusters, among which we found 13 unique to MG3 (**Fig. 4A**), including a strong CD47 signal, a synaptic “don’t eat me” signal, in synaptic remodeling (**Fig. 4B**)^42^.

**Fig. 4.**
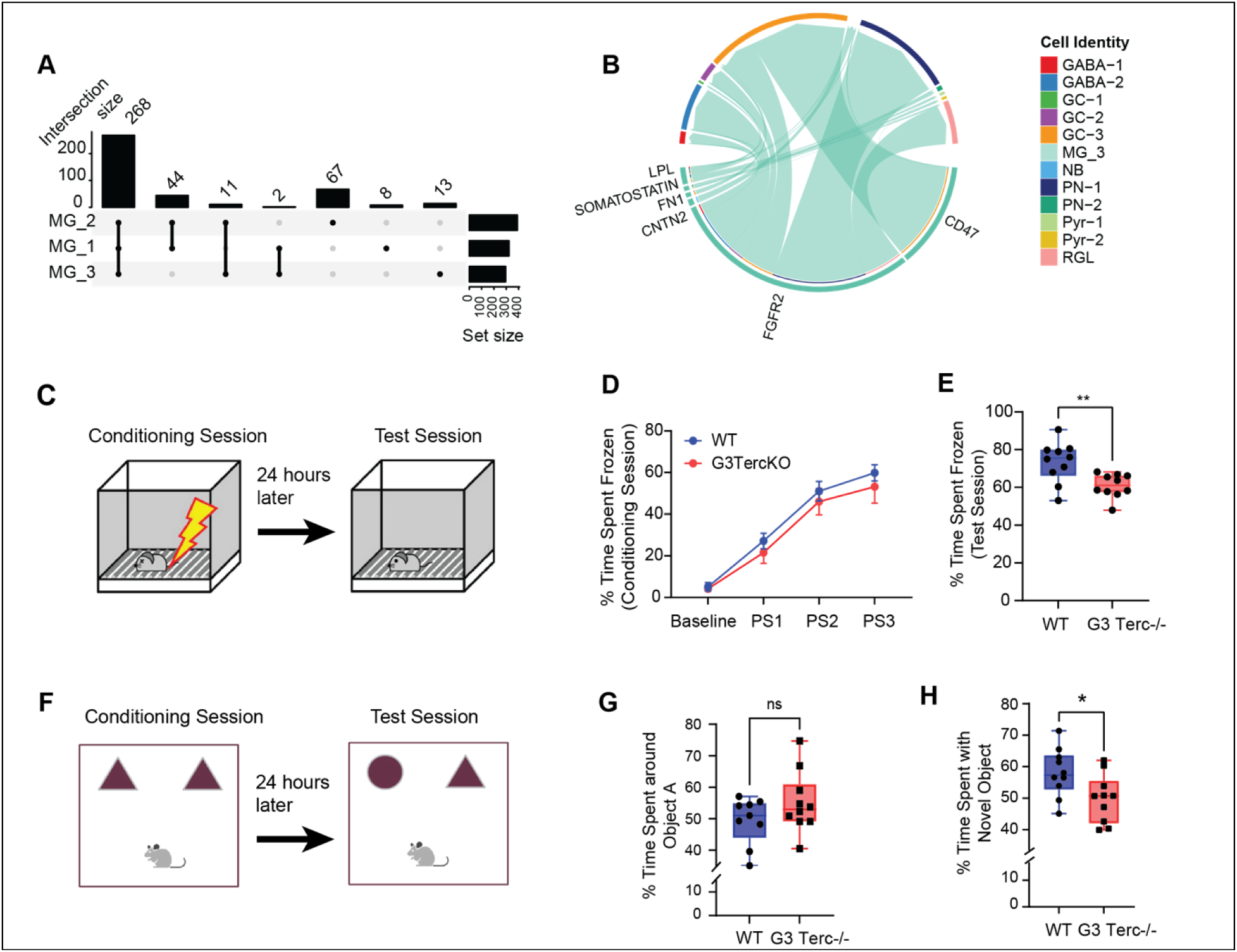
Telomere shortening disrupts microglia-neuron intercellular communication and causes cognitive defect. **A.** Upset plot depicting the relationships of the microglia-subcluster-specific microglia-neuron interactions predicted by CellChat. **B.** Chord diagram showing MG_3-specific signaling pathway predicted by CellChat. The lengths of the segmented outer circle reflect the expression level of ligand proteins in the MG-3 (bottom) and of receptor proteins in the targeted cells (top), showing strong expressions of FGFR2 expression in MG-3 targeting projection neurons and granule cells (GABA: GABAergic neurons; GC: granule cells; NB: neuroblast; PN: projection neuron; Pyr: pyramidal neuron; RGL: radial glial-like cell). **C.** Diagram showing the setup for contextual fear conditioning. **D–E.** Percentage of freezing time the animals spent on the training day (E) and the test day (F). The data show a change in animal behavior at baseline and after receiving shocks. No significant difference was observed on the test day. G3 Terc^-/-^ animals showed a reduction in the percentage time. Each dot represents an animal. Data were reported as box & whisker plot showing min to max and analyzed by unpaired *t*-test. \*\**p*=0.0057. Data were reported as mean ± s.e.m. and analyzed by two-way ANOVA. **F.** Diagram showing the setup for novel object recognition test. **G-H.** Percentage of time animals spent around one of two objects on the training day (H), and around the novel object (I). Each dot represents an animal. Data were reported as box & whisker plot showing min to max and analyzed by unpaired t-test. \**p*=0.0273.

We next determined if telomere shortening caused any age-related deficits on the function level by using contextual fear conditioning (CFC) to assess memory impairment associated with fear (**Fig. 4C**). For CFC, G3 *Terc^-/-^* mice showed no change in the percentage of time frozen during the learning period (**Fig. 4D**). However, the G3 *Terc^-/-^* mice had significantly less frozen time than WT mice during the test period when they were exposed to the same environment without being shocked, suggesting impaired contextual memory (**Fig. 4E**). We then used novel objective recognition (NOR) to test declarative memory (**Fig. 4F**). G3 *Terc^-/-^* and WT mice showed no preference on the two objects as expected during the sample-object exposure session (**Fig. 4G**). In contrast, during the novel-object exposure session, the WT mice showed significant preferences for the novel object, indicating normal memory, but G3 *Terc^-/-^*mice again failed to show a preference for the novel object (**Fig. 4H**). Together, these behavioral deficits indicate that telomere shortening compromises memory function in a manner resembling age related decline^26,43^. Because the manipulation affects all brain cell types, these effects cannot be attributed solely to microglial senescence.

### Soluble DLK1(sDLK1) is identified as a novel senescence and aging-associated microglial ligand

We next sought to determine whether the microglial phenotypes observed in telomere shortened mice could be recapitulated in a human system and to enable causal mechanistic studies using human iPSC-derived microglia like cells (hiMGLs)^44^. We first applied the TERT inhibitor BIBR1532 to iPSCs for 30 days, which significantly reduced telomere length before differentiation into human microglial-like cells quantified with qPCR^45^ (**Fig. 5A,B**). Because hiMGLs are postmitotic, additional telomere shortening after differentiation is unlikely. BIBR1532 treatment did not disrupt microglial differentiation, as indicated by stable expression of canonical microglial markers (**Supp Fig. 3A**). However, telomere dysfunction-induced genes were elevated, consistent with persistent telomere-associated stress, though contributions from non-telomeric DNA damage cannot be excluded (**Supp Fig. 3B**).

**Fig. 5.**
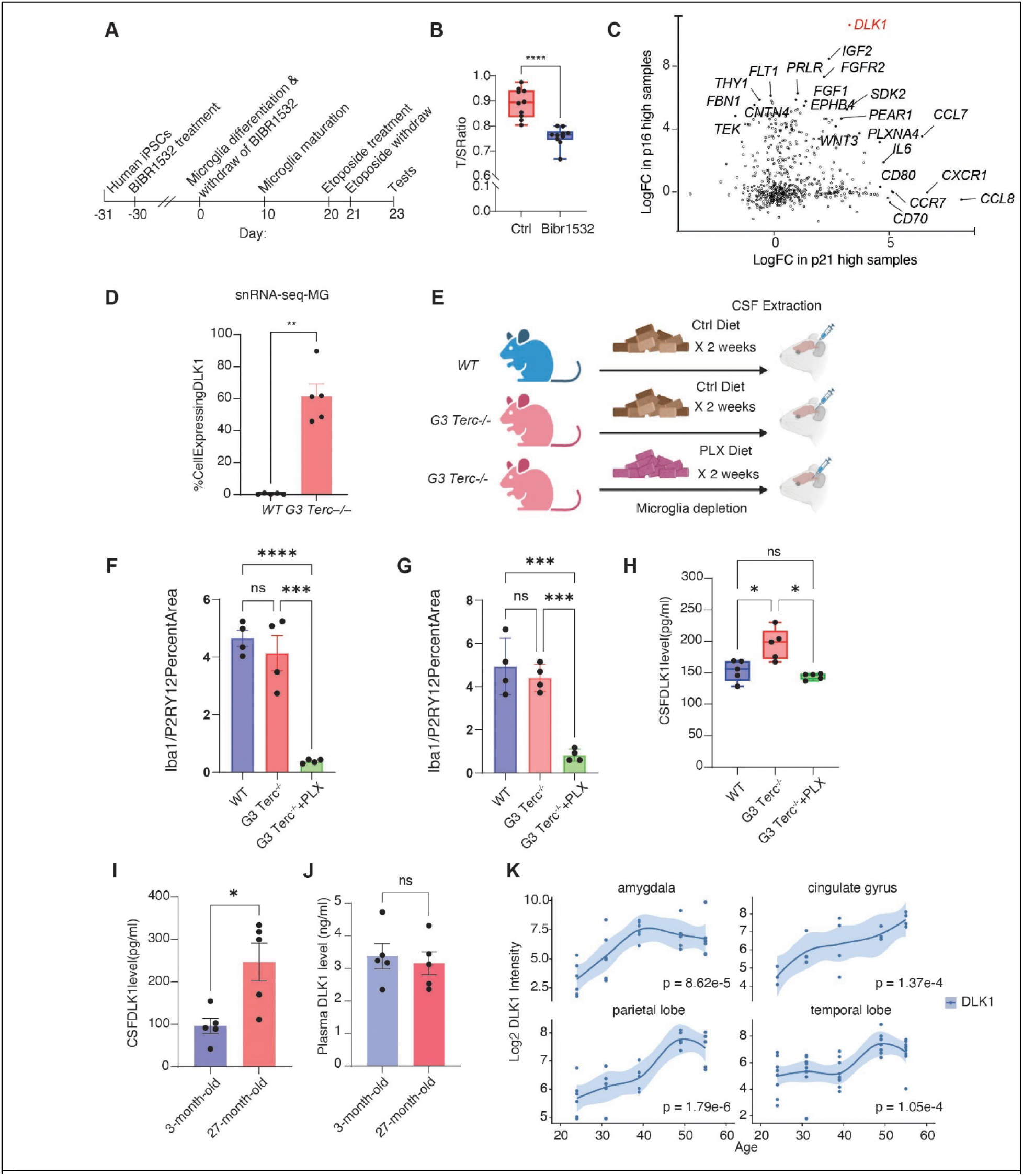
Establishment of a human iPSC-derived microglial senescence model and identification of DLK1 as a microglial-senescence-associated secretory protein. **A.** Schematics of inducing microglia senescence from human iPSCs. **B.** Quantification of iPSC telomere length measured by qPCR. Data were reported as box & whisker plot showing min to max and analyzed by unpaired *t*-test. \*\*\*\**p*<0.0001. **C.** Correlation plot showing the genes correlated to expression levels of p16 and p21 in senescent microglia. DLK1 was strongly correlated with both. **D.** Percentage of microglia expressing DLK1 mRNA in the snRNA dataset. Data are reported as mean ± s.e.m. and analyzed by t-test. \*\**p*=0.0079. **E.** Schematic of microglia depletion experiment set up. The animals were fed normal food or food containing PLX5622 before CSF extraction. **F-G.** Quantification of Iba1+ microglia in CA1 (F) and CA3 (G). 4 mice per condition. 2-3 hippocampal sections/mouse were imaged and analyzed. Data are reported as mean ± SEM. \*\*\*\**p* <0.0001, \*\*\**p* <0.001. Data were analyzed by one-way ANOVA. **H.** Quantification of DLK1 concentration measured in the mouse CSF. G3 Terc^-/-^ mice had elevated DLK1 level in the CSF, and the elevation was diminished by PLX5622 treatment. Data were reported as box & whisker plot showing min to max and were analyzed by one-way ANOVA. (WT vs. G3 Terc^-/-^ * *p*=0.0328) (G3 Terc^-/-^ vs. G3 Terc^-/-^+PLX \**p*=0.0273). Each dot represents an animal. **I-J.** Quantification of DLK1 concentration measured in the CSF (I) or plasma (J) of young (3-month-old) and old (27-month-old) mice. Each dot represents an animal, and the data were analyzed by t-test. \**p*=0.0138. Each dot represents an animal, and the data were analyzed by t-test. **K.** Expression dynamics of DLK1 as a function of age in the human amygdala, cingulate gyrus, parietal lobe, and temporal lobe. p values were obtained by testing the significance of the spline-based smooth term in a generalized additive model.

Senescent cells in vivo typically co-express p16 and p21. To induce p16 and p21 in hiMGLs, we used BIBR1532 pretreatment to reduce telomere length and elevate p16, followed by etoposide to induce additional DNA damage and upregulate p21. The resulting hiMGLs therefore represent a combined telomere erosion plus genotoxic stress phenotype rather than telomere-specific damage alone. Bulk RNA-seq clustered samples into four groups (**Supp Fig. 3C**), confirming the specificity of each treatment: BIBR1532 alone induced p16, etoposide alone induced p21, and only the combined treatment upregulated both markers. Differential expression analysis of the double-positive hiMGLs versus untreated controls identified dysregulated interleukin signaling and secretory pathways (**Supp Fig. 3D**; **Supplementary Table 2**).

We next investigated intercellular signaling changes associated with microglial senescence, focusing on ligand genes curated in CellPhoneDB v2.0^46^. Correlating ligand expression with p16 and p21 identified DLK1, a canonical NOTCH ligand, as the top candidate (**Fig. 5C**). Weighted gene coexpression network analysis (WGCNA) yielded 27 modules, eight of which correlated with DLK1 or the senescence markers (**Supp Fig. 4A, B**). The brown and green modules showed the strongest associations and were enriched for hippo signaling, steroid hormone biosynthesis, and complement pathways, suggesting that DLK1 is integrated into transcriptional programs linked to microglial senescence and immune dysfunction (**Supp Fig. 4C–E**). DLK1 expression in the adult brain is normally restricted to neurogenic niches, where it regulates stem cell activity^47,48^. In contrast, our snRNA-seq analysis revealed low but detectable DLK1 transcripts in more than 60% of senescent G3 *Terc^−/−^* microglia, whereas wild-type microglia lacked DLK1 expression (**Fig. 5D**). This finding supports the idea that DLK1 is aberrantly induced in senescent microglia in vivo.

To test whether senescent microglia in G3 *Terc^−/−^* mice produce and secrete DLK1, we depleted microglia with the CSF1R antagonist PLX5622^49^ (**Fig. 5E**). PLX5622 treatment removed more than 90% of microglia in hippocampal CA1 and CA3, confirmed by loss of Iba1 and P2Y12 staining (**Fig. 5F, G**). We then quantified soluble DLK1 in cerebrospinal fluid by ELISA. Consistent with our snRNA-seq data and hMGL findings, sDLK1 levels were elevated in the CSF of G3 *Terc^−/−^* mice. Importantly, microglial depletion eliminated this increase, demonstrating that senescent microglia are the dominant source of elevated sDLK1 in vivo (**Fig. 5H**).

We next asked whether DLK1 elevation in senescent microglia reflects a broad feature of physiological brain aging. ELISA measurements of CSF and plasma from 3-month-old and 27-month-old mice showed a marked increase in CSF DLK1 in aged animals, with no corresponding change in plasma (**Fig. 5I, J**). These findings indicate that DLK1 elevation is a brain specific aging related alteration rather than a systemic change.

Because DLK1 increases with age in both mouse microglia and our human iPSC iMGLs, we next examined whether similar changes occur in the aging human brain. Using anatomical transcriptomic data from the Allen Institute, we evaluated DLK1 expression across brain regions as a function of age^50^. Several structures, including the amygdala, cingulate gyrus, parietal lobe, and temporal lobe, showed age associated increases in DLK1 expression (**Fig. 5K**).

### sDLK1 disrupts normal functions of oligodendrocyte lineage cells in vivo

Having established that sDLK1 increases with age and is produced by senescent microglia, we next asked how elevated sDLK1 influences brain cell function in vivo. We overexpressed sDLK1 using AAV PHPeB delivered intravenously and confirmed robust induction of sDLK1 in brain tissue two months later (**Fig. 6A, B**). Single nucleus RNA sequencing of the hippocampus identified eight major cell classes using the same quality control criteria applied to the G3 *Terc^−/−^*dataset (**Supp Fig. 5**). sDLK1 transcripts were most elevated in astrocytes and interneurons in AAV-sDLK1 mice (**Supp Fig. 6A**). CellChat analysis showed that astrocytes and interneurons were the major sources of outgoing DLK1 signals, which prominently targeted oligodendrocytes and OPCs (**Supp Fig. 6B**). The strongest predicted interaction was between DLK1 and ErbB4 receptors (**Supp Fig. 6C**), a signaling axis that is essential for oligodendrocyte development and CNS myelination^51,52^. These results pointed to a direct effect of sDLK1 on the oligodendrocyte lineage.

**Fig. 6.**
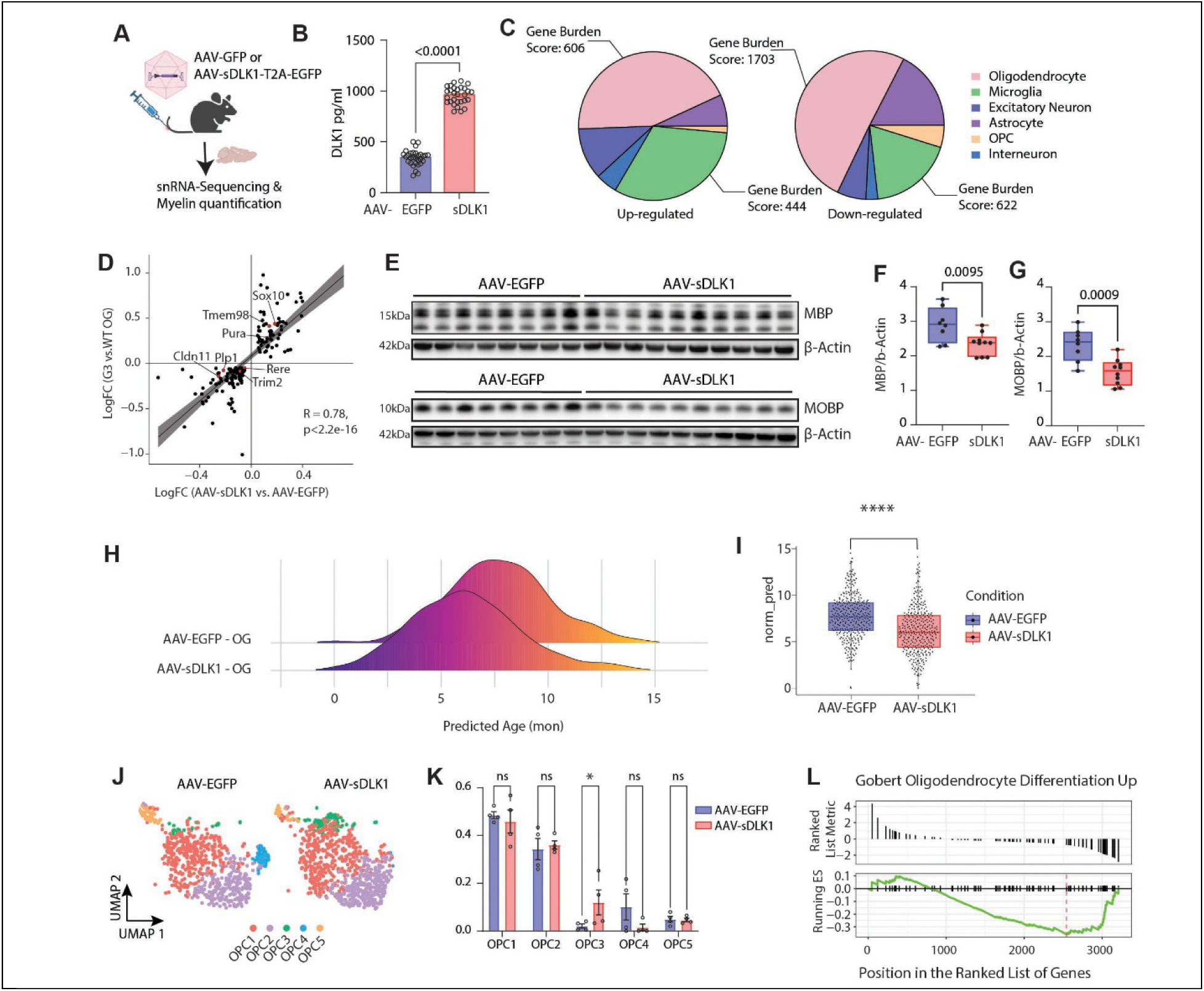
Soluble DLK1 impairs myelination and oligodendrocyte differentiation in adult mice. **A.** Schematic showing the IV injection of the AAV-EGFP or AAV-sDLK1-T2A-GFP into the mice. **B.** Quantification of the protein levels of sDLK1 in the cortex of the mice, measured by mouse DLK1 ELISA. **C.** Distributions of the normalized numbers of genes up-regulated (top) and down-regulated (bottom) in different cell types of the mice injected with AAV-sDLK1-T2A-GFP, compared to mice injected with AAV-EGFP. The gene burden score is defined as the number of differentially expressed genes per 1000 UMI detected in each cell type. **D.** Scatter plot showing the positively correlated genes between DEGs in G3 Terc-/- and AAV-sDLK1-T2A-GFP injected oligodendrocytes. **E.** Western blot of MBP (top) and MOBP (bottom). **F–G.** Quantification of the MBP (F) and MOBP (G) western blot. Data were analyzed by two-tailed unpaired *t*-test. \*\**p*=0.0095 (MBP); \*\*\**p*=0.0009 (MOBP). **H.** Ridge plot of the predicted chronological ages for oligodendrocytes in the mice injected with AAV-EGFP (top) and AAV-sDLK1-T2A-GFP (bottom). **I.** Quantification of the predicted age of oligodendrocytes. Data were reported as a box & whisker plot showing min to max and analyzed by t-test. Each dot represents a cell. \*\*\*\**p* < 0.0001. **J.** UMAP plot showing the enrichment of OPC2 caused by increased sDLK1. **K.** Ratios of each OPC cluster. Data are reported as mean ± s.e.m. and analyzed by two-way ANOVA. \**p*=0.0432. **L.** Running enrichment score and pre-ranked list showing a negative enrichment of oligodendrocyte differentiation predicted by OPC3 markers

Oligodendrocytes exhibited the largest transcriptional response to sDLK1 overexpression (**Fig. 6C**; **Supplementary Table 3**). Although the overlap between DEGs in G3 *Terc^−/−^* mice and AAV-sDLK1 mice is modest (**Supp Fig. 6D**), the shared signals converge on a coherent set of pathways related to myelination and oligodendrocyte maturation (**Fig. 6D**). These common DEGs were selectively enriched for demyelination associated processes (**Supp Fig. 6E**), and 50 of the top 500 were annotated for abnormal myelination or neurological disease (**Supp Fig. 6F**). Together, these focused yet limited shared changes suggest that oligodendrocyte dysfunction in G3 Terc knockout mice is recapitulated in part by sDLK1 exposure, particularly through pathways governing myelin integrity and differentiation.

We next quantified MBP and MOBP and found both significantly reduced in sDLK1 expressing animals (**Fig. 6E-G**). The magnitude of MBP loss within two months suggests active disruption of myelin integrity rather than diminished turnover alone. Importantly, oligodendrocyte and OPC numbers remained unchanged (**Supp Fig. 7**), indicating that sDLK1 impairs myelin maturation and stability without affecting lineage cell abundance in adult mice.

Transcriptomic age prediction further revealed distinct effects of telomere shortening and sDLK1. G3 Terc knockout oligodendrocytes showed higher predicted age, whereas AAV-sDLK1 shifted oligodendrocytes toward transcriptional states characteristic of earlier developmental stages (**Fig. 6H, I**), supporting impaired developmental maturation. Indeed, single nucleus RNA-seq analysis identified five OPC subclusters, with OPC3 enriched in sDLK1 treated brains (**Fig. 6J, K**). Although OPC numbers were unchanged, shifts in OPC states can directly affect myelin quality. OPC3 displayed reduced Tns3, a differentiation associated gene, and increased Grin2b, linked to white matter abnormalities (**Supplementary Table 4**) ^53,54^. GSEA showed that OPC3 markers were negatively enriched for oligodendrocyte differentiation pathways in the sDLK1 treated brain (**Fig. 6L**), indicating that elevated sDLK1 promotes an OPC population with impaired differentiation potential and reduced capacity to support myelin integrity.

### sDLK1 disrupts maturation of human iPSC-derived oligodendrocytes lineage cells

Building on the mouse studies showing that elevated sDLK1 disrupts oligodendrocyte maturation and myelination pathways, we next asked whether sDLK1 exerts similar effects in a human system. We differentiated human iPSCs into oligodendrocyte lineage cells, and treated with recombinant human sDLK1 to mimic the elevation of sDLK1 in the *G3 Terc^−/−^* model (**Fig. 7A**). Single-cell RNA sequencing revealed that the cultures contain both OPCs and mature oligodendrocytes regardless of sDLK1 treatment (**Fig. 7B**). UMAP embedding revealed comparable overall clustering between control and sDLK1 treated cultures, indicating that global lineage structure was maintained, as indicated with expression of PDGFRA, CSPG4, for OPC and MBP for mature oligodendrocyte states (**Fig. 7C–F**).

**Fig. 7.**
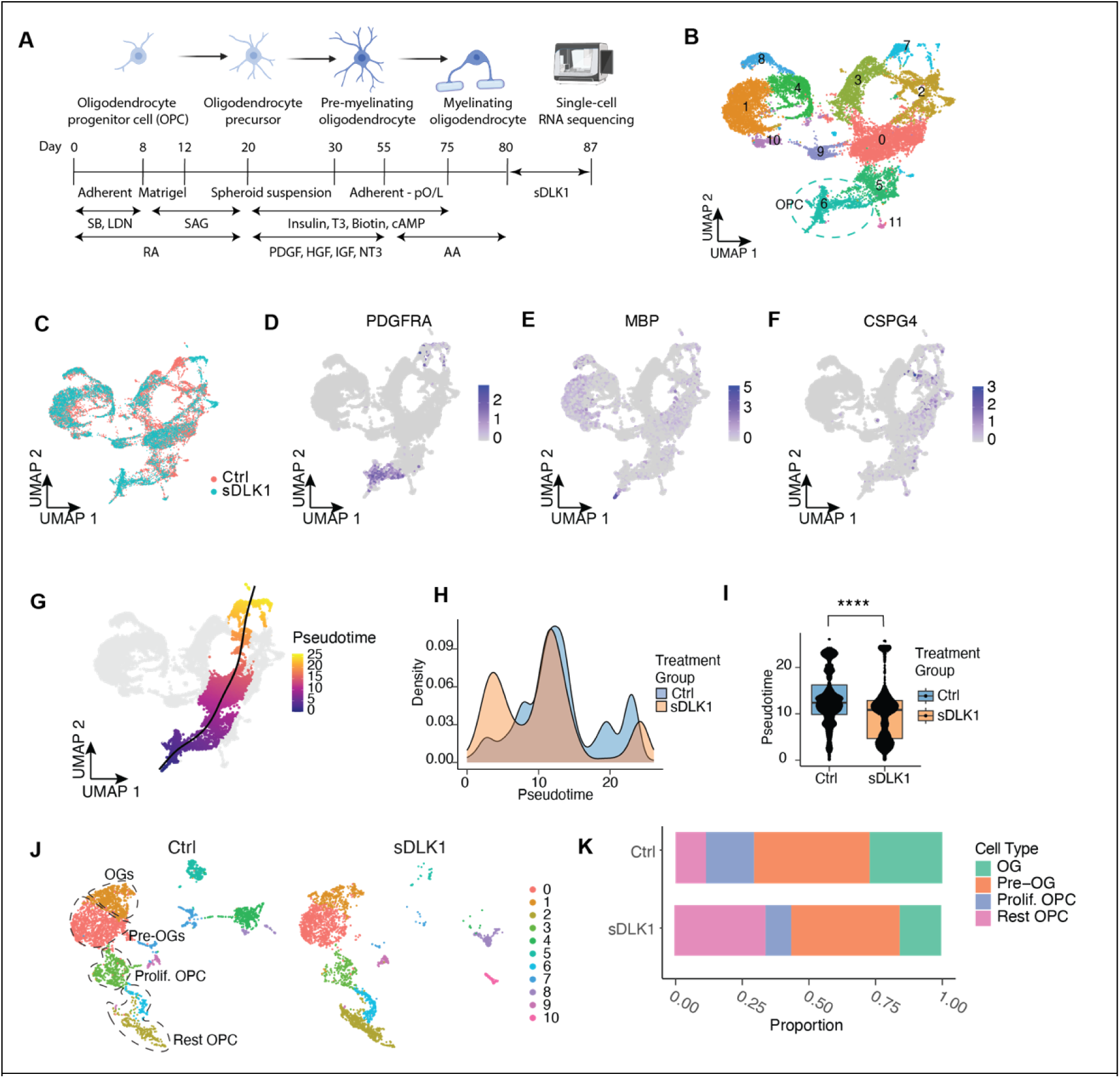
Soluble DLK1 disrupts maturation of human iPSC-derived oligodendrocytes lineage cells. **A.** Schematic showing the differentiation of iOLs from iPSCs. **B.** UMAP plot showing clusters of iOL cultures after removal of neuronal and astrocytic clusters, with cluster 6 defined as the OPC cluster based on PDGFRA expression. **C.** UMAP plots showing iOLs treated with control and recombinant human sDLK1. **D–F.** Feature plot showing the expression of PFGFRA (I), MBP (J), and CSPG4 (K). **G.** Dim plot representing pseudotime lineage originating from OPCs leading to mature oligodendrocyte. **H.** Density plot showing the pseudotime of oligodendrocytes treated with control and sDLK1. **I.** Quantification of the oligodendrocyte pseudotime. Data were reported as a box & whisker plot showing min to max and analyzed by nested *t*-test. Each dot represents a cell. **J.** UMAP plot showing the cells included in the iOL lineage identified by Slingshot. K. Proportion of each oligodendrocyte subtype in the iOL lineage identified by Slingshot.

Using the OPC cluster as a starting point, a Slingshot pseudotime analysis identified an oligodendrocyte maturation lineage and showed that sDLK1 treatment induced a disrupted distribution of cells along the normal progression from OPCs into mature oligodendrocytes (**Fig. 7G, H**). Cultures exposed to sDLK1 experience a significant delay of maturation progress and show an accumulation of cells at the earlier developmental stage as predicted by Slingshot, supporting a block in lineage progression (**Fig. 7I**).

Focusing specifically on the maturation lineage, we identified expected oligodendrocyte subtypes, including pre oligodendrocytes, mature oligodendrocytes, proliferating OPCs, and resting OPCs (**Fig. 7J**). sDLK1 treatment increased the proportion of resting OPCs (**Fig. 7K**), consistent with a block in lineage advancement. Together, these results provide direct evidence that sDLK1 interferes with human oligodendrocyte maturation, mirroring our in vivo findings and establishing sDLK1 as a senescence associated effector that disrupts oligodendrocyte maturation in mouse and human.

### sDLK1 alters calcium transients in human excitatory neurons

We next asked whether DLK1 also perturbs neuronal physiology. Comparing the DEGs in G3 *Terc^-/-^* neurons and AAV-sDLK1-T2A-GFP neurons, we found that 297 of 746 upregulated DEGs in AAV-sDLK1-T2A-GFP neurons were also upregulated in the G3 Terc^-/-^ neurons, demonstrating a remarkable overlap (**Fig. 8A**). Calcium-signal-related pathways were among the top pathways predicted by the DEGs shared between AAV-sDLK1-T2A-GFP and G3 Terc-/- neurons (**Fig. 8B**). GSEA analysis of the overlapping DEGs indicated that, relative to EGFP controls, sDLK1-expressing excitatory neurons showed positive enrichment of calcium ion transmembrane transport and negative enrichment of calcium ion binding, leading us to hypothesize that DLK1 induces abnormal calcium signaling in the excitatory neurons (**Fig. 8C, D**).

**Fig. 8.**
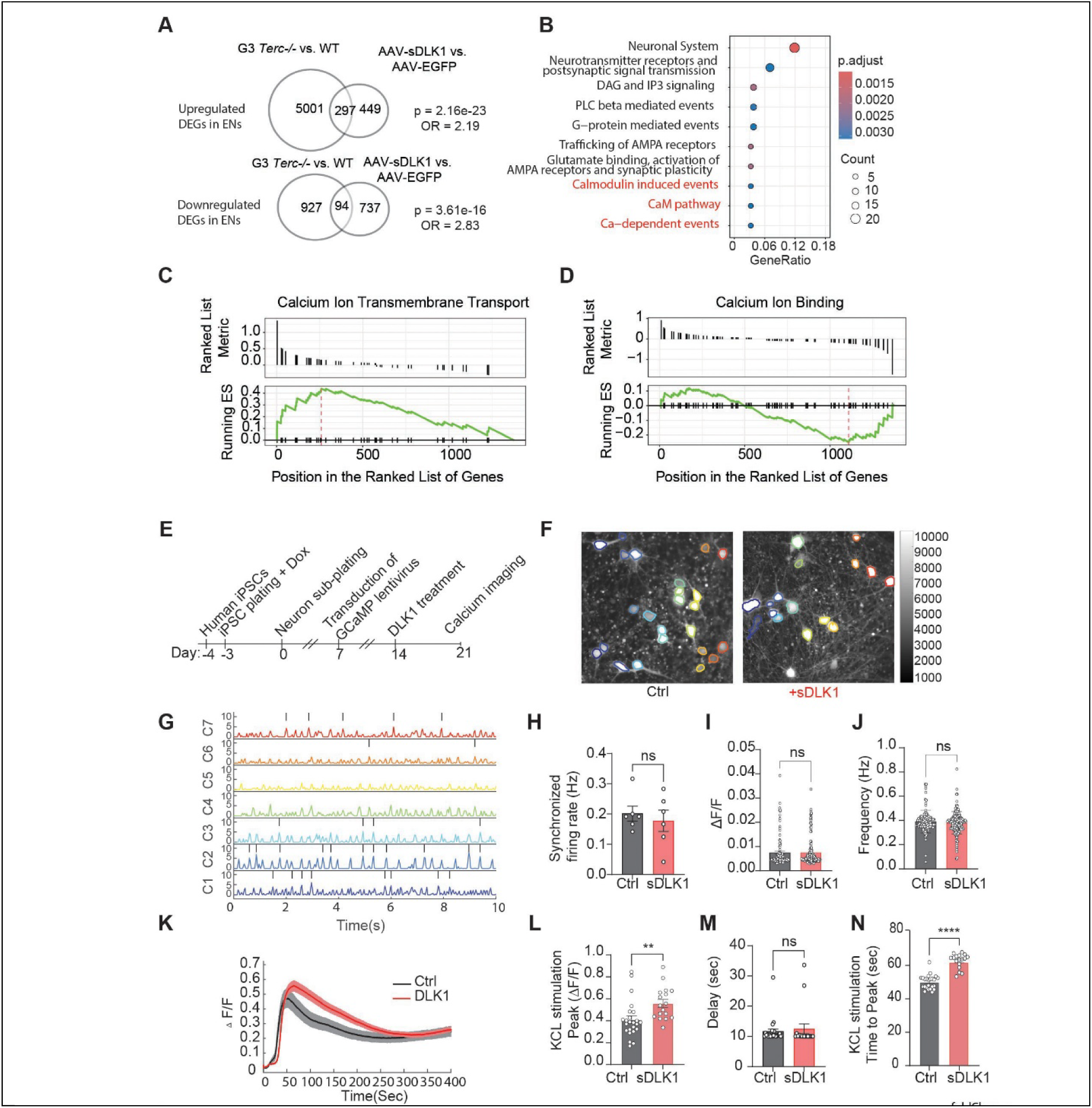
sDLK1 alters calcium transient in human neurons. **A** Venn diagram showing the overlap of upregulated (top) and downregulated (bottom) DEGs in G3 Terc-/- and AAV-sDLK1-T2A-GFP -injected excitatory neurons. **B.** Dot plot of the top 10 Reactome pathways inferred by the upregulated overlapping DEGs in G3 Terc-/- and AAV-sDLK1-T2A-GFP injected excitatory neurons. **C–D.** Running enrichment score and pre-ranked list showing a positive (C) and negative (D) enrichment of calcium ion transmembrane transport predicted by the upregulated overlapping DEGs in G3 Terc-/- and AAV-sDLK1-T2A-GFP injected excitatory neurons. **E.** Schematic illustrating the experiment setup to test the chronic effects of DLK1 on neuronal activities. **F.** Representative fluorescence image of human iPSC-derived neurons expressing GCaMP8f showing spontaneous activity. **G.** Representative spontaneous calcium traces. **H–J.** Quantification of synchronized firing rate (H). firing amplitude (I), or spontaneous firing rate (J). **K.** Representative averaged calcium traces from one KCl stimulation experiment in neurons treated with DLK1 (red) and the untreated control neurons (black). Recording 400 seconds. Mean± s.e.m. **L–N.** Quantification of peak amplitude (L), the delayed KCl stimulation-induced neuronal responses (M), the time each neuron spent to reach peak intensity (N), from KCl stimulation-induced neuronal responses. Data are presented as mean ± s.e.m. and analyzed by unpaired *t*-test. \*\**p*=0.0055 (L) \*\*\*\**p*<0.0001 (N).

We then performed transcriptomic profiling of sDLK1-treated neurons (**Supp Fig. 8A**). Differential expression analysis revealed that most of the affected genes were associated with synaptic compartments, indicating that heightened sDLK1 signaling disrupts synaptic structure and function (**Supp Fig. 8B**). Intriguingly, a subset of sDLK1-induced DEGs overlapped with gene signatures observed in patients at incipient stages of AD^55^, implicating a role of sDLK1 in early synaptic vulnerability and disease progression (**Supp Fig. 8C**).

To determine whether DLK1 alters calcium signaling in human neurons, we expressed the calcium reporter GCaMP8f in human iPSC derived excitatory neurons at day 7 of differentiation and exposed cultures to recombinant human sDLK1 from days 14 to 21 to model sustained ligand elevation (**Fig. 8E**)^56^. Single neuron calcium imaging revealed that chronic sDLK1 treatment did not disrupt spontaneous network activity: synchronized firing rates were unchanged, and the amplitude and frequency of spontaneous calcium transients were preserved (**Fig. 8F–J**). Thus, baseline excitability and network synchrony remain intact. In contrast, sDLK1 significantly altered responses to depolarizing challenge. Upon KCl stimulation, sDLK1 treated neurons exhibited increased peak calcium amplitude (**Fig. 8K, L**). In addition, despite showing no delay in response onset (**Fig. 8M**), neurons treated with sDLK1 required more time to reach maximal signal intensity (**Fig. 8N**),. These changes in evoked calcium dynamics align with the DLK1 induced transcriptional programs and mirror calcium handling deficits described in aged CA1 neurons^57^.

## Discussion

In this study, we demonstrated that telomere shortening, an evolutionarily conserved aging mechanism, causes pathological aging phenotypes within the mouse brain. We comprehensively investigated the underlying mechanisms by which telomere shortening exerts detrimental effects on the brain. Telomere shortening results in a premature accumulation of lipofuscin, a prominent hallmark of aging in the brain. Lipofuscin is a complex mixture primarily of oxidized protein and lipid remnants that resists cellular degradation and elimination and tends to accumulate in post-mitotic cells, such as neurons^26^. Formation of lipofuscin arises from the impaired degradation of damaged mitochondria by lysosomes. Intriguingly, the connection between lipofuscin and telomere shortening was made in a recent study where telomere shortening in the leukocyte was associated with the accumulation of lipofuscin in the serum^58^. Conventionally, the accumulation of lipofuscin is associated with the mitochondrial-lysosomal axis of aging^59^. However, we found that telomere shortening actively facilitates the accrual of lipofuscin, necessitating further investigations into the interplay of telomere and mitochondrial-lysosomal functions. Telomere shortening was also deleterious to memory, thereby corroborating an additional phenotype commonly observed in aged humans and mice. Our *in-situ* hybridization data showed a significant reduction in telomere length in neuronal and glial cells of the G3 Terc^-/-^ animals. Thus, it is imperative to investigate the mechanistic connection between telomere shortening and memory deficit, specifically to determine if the memory deficit is attributed to an inherent neuronal effect or a secondary glial effect.

*Terc^-/-^* animals are characterized by truncated telomeres at the embryonic stage, resulting in shorter telomeres in all somatic cells than WT animals. Consistently, we found heterogeneous transcriptional responses to telomere shortening across all identified brain cell types, but individual cell types had different susceptibilities. We found that excitatory neurons experienced the most pronounced impact, which contradicted our original hypothesis. We had expected neurons to be less susceptible to telomere shortening as the neuronal cells progress into the post-mitotic stage during early development and undergo fewer cell division cycles than glial cells. This finding prompted us to explore the non-cell-autonomous mechanism by which telomere shortening affects the neuronal transcriptome, as glial cells endure more severe telomere attrition stress due to their inherent capability for proliferation. The small overlap between differentially expressed genes in the major glial cell types reveals that telomere shortening induces diverse effects on the glial population. Remarkably, amidst the small overlap of the differentially expressed genes, AC149090.1 was identified as a commonly upregulated gene in microglia, astrocytes, and oligodendrocytes. AC149090.1 was previously identified as a gene upregulated by aging in the hippocampus and neurogenic regions of the brain, and a major contributor to a transcriptome-based aging clock for naturally aged brain cells^30,60^. Although the mechanism by which AC149090.1 is connected to aging remains unclear, our transcriptomic data suggest a connection to telomere shortening and link natural aging in the glial cells to telomere length.

The neuroimmune system, along with its critical constituent microglia, has drawn substantial attention in the field of research focusing on the pathophysiology of neurodegenerative disorders^61^. Impairments and dysregulations of microglial functions have been implicated in neurodegenerative disorders ranging from AD to frontotemporal dementia^62^. Nevertheless, a clear understanding of the underlying factors that drive the initial deterioration of microglia remains elusive. Unlike the other cell types in the CNS, the microglia population derives from microglia progenitors that originated from the yolk sac during early embryonic development^63^. The progenitors undergo extensive replication cycles to achieve the adult microglia population during the embryonic and early postnatal stages^64^. Despite the common belief that microglia are long-lived cells with slow turnover, new evidence has shown that the microglia population undergoes several rounds of proliferation-mediated renewal during the lifetime in both mouse and human^65^. We hypothesized that telomere attrition induced by microglia self-replication, along with its associated replicative senescence, plays a crucial role in disrupting microglial homeostasis and characterized the microglia with shortened telomeres in our telomere shortening model. Our snRNA-seq data showed a striking transformation in the microglia state upon telomere shortening, resulting in the emergence of a distinct subpopulation exhibiting a prominent senescent signature. Senescent microglia exhibited robust upregulation of IL15, a well-established SASP cytokine, and interferon alpha. Both are key hallmarks of senescence^66^. We detected a strong upregulation of other SASP genes and genes known as pathology-associated microglia markers within the senescent microglia population. In parallel, astrocytes in our TERC model also exhibited senescence-associated transcriptomic shifts, indicating that telomere shortening broadly induces glial senescence. Importantly, the transcriptional signatures were accompanied by clear morphological atrophy marked by reduced cell body volume, shorter processes, and fewer branch points. These changes are consistent with microglial dystrophy described in aging brains, supporting the physiological relevance of the telomere-shortening–associated microglial senescence state we observe and its potential as a tool to study microglia in brain aging^36^. The convergence of telomere shortening, microglial senescence, and dystrophic morphology is compatible with telomere-driven DNA damage as the primary initiating event but does not definitively exclude other possibilities.

A growing body of evidence supports the connection between microglial senescence and neurodegenerative diseases^20,67,68^. However, no investigation has examined the intricate transcriptomic changes within senescent microglia and how senescent microglia exert deleterious effects on their surroundings. In our *in vitro* model of microglial senescence, p16 and p21 were upregulated, indicating a mature senescence phenotype. Importantly, the manifestations of senescence, particularly the SASP, exhibit substantial heterogeneity across different cell and tissue types^69,70^. With our *in vitro* model, we gained a comprehensive insight into the transcriptomic alterations within senescent microglia. Given that p21 and p16 represent differential mechanisms in the establishment and sustenance of senescence, we identified genes with positive correlations with either p21 or p16^71^. Our main objective was to elucidate the mechanism by which senescent microglia contribute to brain aging, so we sought to identify genes involved in intercellular signaling among those correlated with p21 and p16. To our surprise, along with cytokines known to be involved in the SASP, we identified that DLK1 expression was strongly correlated with both p21 and p16. Dlk1 is a single-pass transmembrane protein containing a TACE-mediated cleavage site and is a noncanonical member of the Delta-Notch signaling pathway^72^. Dlk1 expression is high during embryonic development but is restricted in adulthood by genomic imprinting^73,74^. The aberrant elevation of DLK1 expression in senescent microglia might be attributed to dysregulation of the imprinted DLK1-DIO3 locus, as observed in human adipose-derived stem cells undergoing replicative senescence, where upregulation of miRNAs and lncRNAs from this locus has been reported^75^. Using a microglia depletion regimen, we confirmed that the expression level of DLK1 is increased in our G3 Terc^-/-^ animals and that the increase was most likely originated from the microglia. Our snRNA-seq data also point to senescent microglia as the main source of DLK1 expression, reinforcing the link between DLK1 expression and microglial senescence. With the establishment of DLK1 as a novel member of the microglial SASP, we also observed elevated DLK1 levels in the CSF of physiologically aged animals, suggesting a link between DLK1 and aging. Interestingly, the age-related elevation in DLK1 was not observed in peripheral plasma samples, indicating that excessive DLK1 level is a phenomenon specific to brain aging. Transcriptomic data from human postmortem brains further suggest that DLK1 may play a conserved role in brain aging across species.

The involvement of oligodendrocyte lineage cells in brain aging and the development of age-dependent neurodegenerative disorders, including diseases not primarily associated with myelination defects (e.g., AD and PD), has attracted increasing attention in recent years^76^. Our data suggest that elevated DLK1 signaling from senescent microglia disrupts the differentiation and myelinating capacity of oligodendrocytes and likely represents one of several pathways through which telomere shortening impairs white matter integrity. This is consistent with previous findings in which RHEB-knockout-induced upregulation of DLK1 from neurons impaired oligodendrocyte differentiation and myelination^77^. Indeed, we observed loss of myelination and disrupted oligodendrocyte differentiation in the G3 Terc^-/-^ mouse brains. However, the proliferative nature of OPCs renders it inconclusive whether the myelination phenotype in the G3 Terc^-/-^ mouse is attributed to increased DLK1 signaling or to the cell-autonomous effect of telomere shortening. After overexpressing the soluble form of DLK1 in the mouse brain, we confirmed that elevated soluble DLK1 is sufficient to cause defects in myelination and oligodendrocyte differentiation. The OPC3 cluster induced by elevated sDLK1 represents a functionally distinct adult OPC population that retains developmental potential yet fails to complete maturation, clarifying how age-related signaling and SASP factors create remediable barriers to oligodendrocyte differentiation. Furthermore, human iPSC-derived oligodendrocyte cultures exposed to human recombinant sDLK1 exhibited a marked delay in maturation, with cells stalling at precursor stages. In parallel, soluble DLK1 overexpression in vivo perturbed calcium signaling pathways in neurons, which provides a potential explanation for the memory deficits in the G3 Terc^-/-^ animals. Our *in vitro* model confirmed that neurons exposed to human recombinant sDLK1 displayed dysregulated calcium dynamics, including exaggerated intracellular Ca^2+^ influx upon depolarization. Chronic DLK1 exposure also disrupted the expression of synaptic and AD-associated genes, suggesting a mechanistic link between microglial senescence and neuronal vulnerability. These sDLK1-driven alterations in normal oligodendrocyte function and neuronal calcium homeostasis provide mechanistic evidence that senescent microglia can contribute to neurodegenerative pathologies in other cell types through a unique secretory profile.

The mechanisms underlying pathological brain aging and associated neurodegenerative disorders remain largely elusive, especially their involvement in disease development before disease manifestations, such as misfolded proteins. Although telomere length is only one of many metrics of aging, we found that telomere shortening and microglia senescence induced by it can give rise to key phenotypes of pathological brain aging, proving its potential as a tool to further understand the alterations caused by age and early disease development.

While this study provides novel insights into microglial senescence and the role of DLK1, it has several limitations. Our in vivo and in vitro experiments demonstrate that elevated sDLK1 impairs oligodendrocyte maturation, disrupts myelination, and alters neuronal activity, but we did not assess whether these cellular and circuit-level disturbances translate into broader aging-related outcomes, such as cognitive decline, systemic senescence, or activation of senescence-associated secretory programs. Future longitudinal behavioral studies and transcriptomic profiling will be required to clarify whether sDLK1 contributes to higher-order aging phenotypes. We also found that astrocytes and other glial subtypes exhibited senescence signatures, suggesting that additional cell types may contribute to aging-related dysfunction in ways not captured by our current design. In addition, our human iPSC-derived model relied on sequential BIBR1532 and 50 µM etoposide treatment to elicit complementary p16 and p21 pathways, which likely combine effects of telomere shortening with widespread etoposide-induced DNA damage and therefore may introduce confounding effects, precluding a clear separation of telomere-specific versus generalized genotoxic contributions and not fully recapitulating telomere-driven senescence in vivo. Future work using genetic or conditional approaches that perturb telomere maintenance without exogenous genotoxic agents will be important to sharpen causal inferences about telomere-driven senescence. Finally, although we demonstrated a potential mechanism by which telomere shortening causes age-related pathological phenotypes, we did not focus on any disease-specific phenotypes. Another logical follow-up is to combine the telomere shortening model with disease-specific risk genes, such as APOE or TREM2, to further examine the mechanisms underlying the age dependency of neurodegenerative disease and potential methods for early disease diagnosis and intervention.

## SUPPLEMENTARY MATERIALS

### Method

#### Mice

Male and female G0 TERC^-/-^ breeders were purchased from the Jackson Laboratory (The Jackson Laboratory, 004132) and bred for three generations to generate G3 TERC^-/-^ animals. Age-matched WT C57BL6/J mice were obtained from the NIA. Animals were housed no more than five per cage in a pathogen-free barrier facility at 21–23°C with 30–70% humidity on a 12-hour light/dark cycle. The animals were given *ad libitum* access to food and water. Both male and female animals were used for histological analysis. Only female animals were used for behavioral and biochemical analyses. Animals were 8–9 months of age when used for histological analysis. Animals underwent behavioral testing at 9 months of age and had not been used previously for any other experiments. At 10 months of age, the same mice were used for pathology and RNA-seq studies. For microglia depletion regimen, PLX5622 was given to the animals with food at 8 months of age for 2 weeks before the behavioral experiment and the subsequent pathology studies. All mouse protocols were approved by the Institutional Animal Care and Use Committee, University of California, San Francisco, and Weill Cornell Medicine.

#### Brian tissue collection

Mice were euthanized using Fatal-Plus (pentobarbital sodium) and transcardially perfused with PBS. The brains were hemisected, and the hemibrains were flashed-frozen at -80°C or fixed in 4% paraformaldehyde for 48 hours, which was then followed by 48-hour 30% sucrose infiltration at 4°C. The fixed hemibrains were sectioned into 30-µm slices using freezing microtome (Leica) and stored at -20°C in cryoprotectant before staining.

#### Contextual fear conditioning

Mice were tested for contextual fear learning and memory using sound-attenuated chambers (Med Associates, VT, USA). During fear acquisition, mice freely explored a novel environment. After a 2-min habituation, mice were exposed to a 2-s foot shock (0.5 mA) followed by a 60-s interstimulus interval for a total of three shocks. At 24 hours after fear acquisition, hippocampal-dependent fear memory was measured by recording the percent total time spent freezing in a 5-min context test (Video Freeze software).

#### Novel object recognition

Mice were habituated to opaque open field arenas (40 × 40 cm) by allowing them to freely explore the arena for two 10-min trials spaced on the 2 days leading up to the object recognition test. On the test day, two identical objects (plastic geometric object) were placed in the center of each arena. Mice were allowed to freely explore the objects for a 15-min trial. After 24 hours, one of the geometric objects was replaced by a novel object of a different shape and color. The animal was then allowed to freely explore the new objects for a 15-min trial. Video recording and tracking (Ethovision v15, Noldus) were used to track the movement of animals. The time mice spent exploring each object was determined automatically by the Ethovision software. Preference was calculated based on the total time an individual mouse spent exploring both objects.

#### Telomere length measurement

Telomere length labeling was done using TelC-Cy3 (PNA Bio Cat. No. F1002), according to the manufacturer’s protocol with modifications. PFA-fixed brain sections were incubated in 1% Tween-20 in PBS for 1 minute, followed by boiling in antigen unmasking solution (Vector Cat. No. H-3300) at 90 °C for 35 minutes. After cooling for 5 minutes, the sections were rinsed with PBS. In a PCR tube, 50 μL of TelC-Cy3 solution (1:250 dilution of 250 μg/ml formamide TelC-Cy3 stock solution into PNA staining solution (70% formamide, 10 mM Tris pH 7.5, 0.5% B/M Blocking Reagent solution (Sigma Cat. No. 11096176001, prepared according to manufacturer’s instructions)) was added to a brain section and heated at 84 °C for 5 minutes. The sections were left overnight at room temperature and protected from light. Thereafter, the sections were washed twice for 15 minutes with PNA Wash Solution (70% formamide, 10 mM Tris pH 7.5). Finally, the sections were washed with 0.01% Tween-20 in PBS. To identify TelC-Cy3 signal in microglia, Iba-1 labeling was performed according to a published procedure, and counterstained with DAPI^78^. Brain sections were mounted and imaged using Zeiss LSM880 inverted scanning confocal microscope (Carl Zeiss Microscopy, Thornwood, New York) with 10 series of 1 μm sections. Z-max projections were analyzed; TelC-Cy3 signal was quantified using a published method^79^. Briefly, using FIJI^80^, the background-corrected TelC-C3 signal in the nucleus of Iba-1+ microglia was calculated by subtracting background from the average of top 20% of red pixels. For each genotype, four male and four female mice were analyzed, with three images taken per animal, and 9–20 Iba1+ cells per animal were quantified. On average, WT animals were 8.5 months old, and G3 animals were 8.62 months old.

#### Nuclei isolation from frozen mouse hippocampi

Five female animals of each genotype were used for snRNA-seq. Nuclei isolation was done as described, with modifications^81,82^. In brief, mouse hippocampi were dissected from frozen brain tissue before placed in 1,500 µL of Sigma nuclei PURE lysis buffer (Sigma, NUC201-1KT). Hippocampal samples were homogenized with a Dounce tissue grinder (Sigma, D8938-1SET) with 20 strokes using pestle A, followed by 15 strokes using pestle B. After homogenization, the tissue was filtered through a 35-µm cell strainer, followed by centrifugation at 600g for 5 minutes at 4 °C. The resulting pellet was washed three times with 1 mL of PBS containing 1% bovine serum albumin (BSA), 20 mM DTT, and 0.2 UµL^-1^ recombinant RNase inhibitor. The nuclei sample were then centrifuged at 600g for 5 minutes at 4 °C and subsequently resuspended in 800 µL of PBS containing 0.04% BSA and 1X DAPI, followed by fluorescence-activated cell sorting (FACS) to remove cell debris.

#### Droplet-based snRNA-seq

snRNA-seq libraries were prepared with Chromium Single Cell 3’ Reagent kits (v3; 10X Genomics, PN-1000075). The libraries were sequenced on a NovaSeq 6000 sequencer (Illumina) with 100 cycles.

#### Analysis of droplet-based snRNA-seq data

RNA reads sequenced from the snRNA-seq library were aligned to mm10 genome using Cell Ranger software (v.3.1.0; 10X Genomics) to generate raw gene counts. Reads mapped to pre-mRNA were counted to include un-spliced nuclear transcripts. Cell barcodes were called using Cell Ranger 3.1.0 default parameters. We further removed genes expressed in no more than three cells, cells with unique gene counts over 8,000 or less than 100, and cells with more than 5% mitochondrial reads. High-confidence doublets were removed from individual samples using DoubletFinder^28^. Sample integration, normalization and clustering were done with the Seurat package v3.0.1^83^. In brief, integration anchors were computed for the datasets using the top 30 principal components, and the datasets were integrated using the anchor set. The integrated dataset was scaled by the total library size multiplied by a scale factor (10,000) and transformed to log space. Principle component analysis was done on the highly variable genes, and t-distributed stochastic neighbor embedding was run on the top 30 principal components. k.param was computed using the top 30 principal components, and cells clusters were identified using the FindCluster function with a resolution of 0.02. Uniform Manifold Approximation and Projection (UMAP) was run on the top 30 principal components. Cell-type labels were assigned to teach cluster using statistical enrichment for sets of marker genes and performing manual evaluation of gene expression for small sets of known marker genes. The dataset was then split into individual datasets based on cell-type identity. Differential gene expression analysis was done using the FindMarkers function with MAST as the method used. Ingenuity Pathway Analysis (IPA) and MSigDB gene annotation database were used to identify gene ontology and pathways enriched in the differentially expressed genes (DEGs). To address multiple testing, Benjamini-Hochberg approach was used to generate corrected false discovery rate (FDR). For trajectory analysis, the dataset was converted to a monocle3 object and analyzed using the Monocle3 package^84,85^. Pseudotime analysis was done on the microglia data using WT microglia cluster as the origin. To evaluate cellular interactions between microglia and neuronal cells, we applied CellChat (v.1.6.1) to examine the ligand-receptor interactions inferred by the dataset^86^. Normalized integrated data was used as input, and the analysis followed the CellChat tutorial with default parameters and CellChatDB.mouse as the interaction database. Ligand-receptor interactions were plotted using netVisual_chord_gene and netVisual_bubble functions.

#### Machine learning model to predict cell age using snRNA-seq data

Biological age prediction models were trained using the publicly available single-cell RNA-seq data from the SVZ neurogenic niche, with the code described in a previously published paper^30^. The single-nuclei data of astrocyte, microglia, and oligodendrocyte were preprocessed by the “BootstrapCell” method before being applied to the trained models. The distribution of predicted biological ages was visualized using ridge plots and dot plots.

#### Bulk RNA-seq

Primary microglia and neurons were disassociated from the tissue plates using Accutase (ThermoFisher, 00-4555) and collected by 3-minute centrifugation at 500 × g. mRNA was extracted according to the manufacturer’s protocol (ZYMO Research, *Quick*-RNA Microprep Kit). Isolated RNA was sent to Novogene Co. for quality control, library preparation, and sequencing. RNA-seq read mapping was performed using the STAR program and with the GENCODE GRCh38.p13 as reference. The read count table was generated with the Rsubread package. Differential gene expression was calculated with the DEseq2 package.

#### Lipofuscin imaging and quantification

PFA-fixed hemibrain sections were washed with 0.5% Triton in PBS and mounted for imaging on Keyence BZ-9000 inverted epifluorescence microscope (Keyence, Osaka, Japan). The entire hemibrain section was scanned at 10x magnification using Cy3 channel, and the individual images were stitched together using Keyence BZ-X Analyzer software. All images were thresholded uniformly, and a 400x400 pixel square was cropped in primary somatosensory cortical region for endogenous fluorescence quantification. Using “Analyze Particles” in FIJI, the area of lipofuscin particles was quantified. Four mice for WT (two males and two females, average age: 9 months old) and three for G3 (one male and two females, average age: 9.3 months old) were quantified, with eight sections per mouse. Average numbers of puncta were 93 per animal for Terc^+/+^ and 168 for G3 Terc^-/-^.

#### Cerebrospinal fluid extraction

CSF extraction procedure was described in detail^87^. Briefly, mice were deeply anesthetized using Fatal-Plus (pentobarbital sodium) and positioned prone over a 15-mL conical tube to place the cervical spine in flexion. The mouse occiput was palpated to locate the cisternal magna, and a 30G insulin needle was punctured and advanced less than 4mm deep. CSF was collected by slowly pulling the syringe plunge. Approximately 15 uL CSF was collected from each animal. CSF was spun down at 600g for 5 minutes in 4⁰C. Supernatant was collected, flash-frozen on dry ice and stored at -80⁰C until analysis.

#### ELISA and multiplex bead-based immunoassay

Extracted CSF was diluted in PBS and assayed using the Mouse Pref-1/DLK-1/FA1 ELISA Kit (Invitrogen, EM66RB), according to the manufacturer’s instructions. Other SASP cytokines were measured with a MILLIPLEX MAO mouse cytokine/chemokine magnetic bead kit (Millipore) using a MAGPIX system.

#### Human iPSC differentiation into microglia and senescence induction

iPSCs from an adult female with no known diseases were purchased from WiCell (UCSD072i-1-3) for the differentiation of microglia. Cells were cultured in an incubator at 37°C with 5% CO_2_. iPSCs were passed three times for expansion purpose before the start of experiments. Bibr1532 (60 µM; Millipore Sigma, 508839) was added to iPSCs culture after starting the experiments, and the iPSCs were cultured in BIBr1532-containing medium for 30 days before microglia differentiation. iPSCs were passed at a ratio of 1:12 every time the culture reaches 80% confluency. Bibr1532 was withdrawn after 30 days, and the iPSCs were differentiated into macrophage progenitors via a 10-day protocol, as described^44^. The macrophage progenitors were then further matured into microglia-like cells via a 13-day protocol with macrophage colony–stimulating factor and interleukin 34 in RPMI supplemented with 10% FBS as described^44^. Differentiation quality control was conducted through IBA1 (Abcam, ab5076) and TMEM119 (Sigma, HPA051870) immunocytochemistry. Matured microglia were treated with 50 µM Etoposide (Millipore Sigma, E1383) for 24 hours and then recovered for 24 hours before subsequent experiments.

#### Human iNeuron differentiation

Human iNeurons are differentiated as described, where WTC11, iPSCs from a male, were engineered for inducible expression of Ngn2 from a transgene integrated in the AAVS1 locus^88,89^. The differentiation process followed the published protocol^88^. Briefly, the iPSCs were pre-differentiated to neuronal precursor cells in pre-differential medium containing doxycycline. On day 0, neuronal precursor cells were replated in maturation medium containing doxycycline. Doxycycline was removed from the maturation medium on day 7. Thereafter, one-half of the maturation medium was replaced with fresh medium weekly until the cells were collected.

#### Immunocytochemistry

iPSC-derived microglia and neurons on coverslips were fixed in 4% paraformaldehyde in PBS for 30 minutes and then washed three times for 5 minutes each with PBS. The cells were then permeabilized with 0.1% Triton X-100 in PBS (PBS-T) before blocking with 5% normal donkey serum (NDS) in PBS-T for 1 hour at room temperature. Cells were washed three times with PBS after blocking. Primary antibodies were added in PBS-T and incubated at 4°C overnight. The secondary antibodies were added to the cells for 2 hours at room temperature after washing with PBS for three times. DAPI was added to the cells for nuclei labeling for 10 minutes before visualization. Images were acquired with a laser scanning confocal microscope (Zeiss, LSM 700) using a 63X oil objective or a Apotome3 microscope (Zeiss, Axio Observer). The image acquisition settings were chosen to prevent most of the brightest pixel intensities from reaching saturation. The anatomically uniform corpus callosum was selected for oligodendrocyte staining to allow unambiguous identification and accurate quantification of ANLN-positive oligodendrocytes.

#### Calcium imaging

iPSCs were differentiated into neurons on coverslips as described. Neurons were transduced with hSyn-jGCaMP8f lentivirus on D6. The lentiviral construct was made using the third generation lentiviral plasmid FUGW (Addgene, 14883), where the Pacl + EcoRI fragment was replaced by the hSyn-GCaMP8f fragment^90^. The medium containing the lentiviral construct was replaced by fresh medium on Day 7. DLK1 treatment started on Day 14 (200ng/ml, R&D System, 1144-PR), and the cells were imaged and sequenced on Day 21. At the time of imaging, the coverslip was gently washed and placed into a glass-bottom chamber (Warner Instruments, RC-26G) containing Ca^2+^ imaging buffer (20 mM HEPES, 119 mM NaCl, 5 mM KCl, 2 mM MgCl_2_, 30 mM glucose, 2 mM CaCl_2_, pH 7.2–7.4). The temperature was maintained at 37°C by a dual chamber heat controller (Warner Instruments, TC-344C). Fluorescence time-lapse images were collected on a microscope (Nikon, FN1) using a 60X, 1.0 NA objective (Nikon, CFI APO 60XW NIR) and a C-FL GFP filter cube. An X-CITE LED illuminator (Nikon) was used for excitation. Images were collected using an ORCA-Fusion CMOS camera (Hamamatsu) with 4×4 binning (576×576-pixel resolution, 16-bit grayscale depth, 0.43 µm/pix) and NIS-Elements AR software (Nikon). Exposure time was set to 20 milliseconds. For spontaneous activities, eight fields per coverslip were acquired at 30 Hz for 2 minutes. After imaging spontaneous activity, one filed per coverslip was imaged for 10 minutes at 20 Hz to minimize photo-bleaching immediately after 50 mM KCl perfusion.

#### Generation of AAV

In this study, the pAAV-CAG-GFP plasmid (Addgene, 37825) was used. The extracellular domain of mouse Dlk1 (amino acid residues 1–170; sDLK1), followed by a T2A sequence, was inserted upstream of the GFP gene to generate pAAV-CAG-sDLK-T2A-GFP-WPRE-SV40polyA. AAV-PHP.eB virus was produced at the University of Pennsylvania (https://gtp.med.upenn.edu/vector-core). Intravenous injections of AAV particles encoding mouse sDLK1 were performed in C57BL/6 mice, while PHP.eB AAV encoding GFP alone (Addgene, 37825-PHP.eB) served as a negative control. Each AAV preparation had a titer of 1 × 10¹¹ viral genomes per mouse, and both male and female mice were included. One to two months after injection, mice were anesthetized and perfused with ice-cold PBS, and samples were harvested for biochemical or histological analyses.

#### Western Blot

For mouse brain samples, half cortex was mechanically homogenized on ice in RIPA buffer containing protease and phosphatase inhibitors (Millipore Sigma). Fifty micrograms of cortex lysates were loaded onto NuPage Bis-Tris gels (Thermo Fisher) and run in SDS running buffer at 150 V for ∼2.5 h. Gels were transferred to nitrocellulose membranes (Bio-Rad) overnight in a cold room. Membranes were washed three times for 10 min each in TBS with 0.01% Triton X-100 (TBST) and blocked for 1 h in 5% milk in TBST. Appropriate primary antibodies were diluted in 1% milk in TBST and incubated at 4 °C overnight. The following day, membranes were washed three times for 10 min each in TBST and incubated with appropriate secondary antibodies in 1% milk in TBST for 1 h at room temperature. Membranes were washed again to minimize nonspecific binding, treated with ECL (Bio-Rad) for 60 s, and developed in a dark room or using a Bio-Rad imager. Blots were scanned at 300 d.p.i. and quantified using ImageJ.

#### Antibodies

Antibodies used in immunofluorescence analyses were as follows. Secondary antibodies used were Alexa Fluor donkey anti-rabbit/goat 488 and anti-mouse 555 (Invitrogen) and donkey anti-goat 555 at a dilution of 1:500 (Jackson ImmunoResearch). Primary antibodies included anti-ANLN (ORIGENE, TA349469, 1:200), anti-OLIG2 (Millipore, MABN50, 1:500), anti-MBP (Millipore, MAB386, 1:1000), anti-MOBP (Thermo Fisher Scientific, AB-2718347, 1:500), anti-Iba1 (1:500, Abcam, cat. # ab5076), anti-P2RY12 (1:250, BioLegend cat. # S16007D), anti-PDGFRα (1:500, R&D Systems cat. # AF1062). Secondaries used for western blotting were anti-rabbit horseradish peroxidase (Calbiochem, 401393; 1:2,000) or anti-mouse horseradish peroxidase (Calbiochem, 401253; 1:2,000).

#### qPCR-based telomere length measurement

Relative telomere length was measured in iPSCs by real-time qPCR as described in a previous study^91^. Briefly, genomic DNA was extracted from iPSCs using PureLink^TM^ Genomic DNA Mini Kit (ThermoFisher, K182002). Same amount of genomic DNA was loaded onto a 96-well plate with telomere amplification primers (Telg: ACACTAAGGTTTGGGTTTGGGTTTGGGTTTGGGTTAGTGT & Telc: TGTTAGGTATCCCTATCCC-TATCCCTATCCCTATCCCTAACA) and single gene (β-globin) amplification primers (Hbgu: GCTTCT-GACACAACTGTGTTCACTAGC & Hbgd: CACCAACTTCATCCACGTTCACC). A standard curve was generated by measuring the Ct (cycle threshold) at which fluorescence exceeds a defined baseline and plotting Ct versus log(input DNA concentration) from serial dilutions of a reference sample. Telomere (T) and single-copy gene (S) assays were run on the same 96-well plate and T/S ratios for unknowns were determined by interpolating their Ct values against the standard curve, yielding precise relative telomere-length measurements.

#### Microglia Morphology Analysis

63x confocal z-stack images (0.44 µm step size, 0.13 µm pixel size) were acquired on a Zeiss LSM 880 microscope. 3-4 sections/mouse were imaged in the CA1 region of the hippocampus and 4-9 fields of view were imaged/section. Prior to morphological analysis, images underwent smoothing and background subtraction in Imagej/Fiji. Microglia were then surface rendered using Imaris software and microglia volume was calculated. Microglia surfaces were then masked and traced using Filament Mode in Imaris to calculate the total process length/microglia and the number of branch points/microglia.

#### OPC/OL Differentiation from iPSCs

The human iPSC line WTC11 was differentiated into oligodendrocytes following a published protocol^92^. Briefly, iPSCs were dissociated into single cells using Accutase (Sigma-Aldrich, Cat# A6964) and plated at a density of 1×10^5 cells per well on Matrigel-coated 6-well plates. From day 0–8, cells were cultured in neural induction medium consisting of mTeSR™ Plus (Stem Cell Technologies, Cat# 100-0276) supplemented with 1× GlutaMAX, 1× NEAA, 10 μM SB431542 (Tocris, Cat# 1614), 250 nM LDN193189 (Sigma, Cat# SML0559), and 100 nM retinoic acid (Sigma, Cat# R2625). From day 8–12, cells were maintained in N2 medium (DMEM/F12 supplemented with 1× N2, 1× NEAA, 1× GlutaMAX, 100 nM retinoic acid, and 1 μM SAG (Millipore Sigma, Cat# 566660)). From day 12 to day 20, cells were lifted into spheroids using cell scrapers and cultured in N2B27 medium (N2 medium supplemented with 1× B27 minus vitamin A). From day 20–29, spheroids were cultured in PDGF medium (DMEM/F12, 1× N2, 1× B27 minus vitamin A, 1× NEAA, 2 mM GlutaMAX, 10 ng/mL PDGF-AA (R&D Systems, Cat# 221-AA-050), 10 ng/mL IGF-1 (R&D Systems, Cat# 291-G1-200), 5 ng/mL HGF (R&D Systems, Cat# 294-HG-025), 10 ng/mL NT3 (Millipore, Cat# GF031), 60 ng/mL T3 (Sigma-Aldrich, Cat# T2877), 100 ng/mL biotin (Sigma-Aldrich, Cat# 4639), 1 μM cAMP (Sigma-Aldrich, Cat# D0260), and 25 μg/mL insulin (Sigma-Aldrich, Cat# I9278)). From day 30 onward, spheroids were plated on poly-L-ornithine/laminin–coated dishes and maintained in PDGF medium to promote OPC generation and further oligodendrocyte maturation.

#### Single-cell RNA-seq of OPC/OL Differentiation from iPSCs

After dissociation, cells were suspended in 1× PBS with 0.04% BSA for a final concentration of 1,000 cells µl^−1^. scRNA-seq libraries were prepared with Chromium Single Cell 3’ Reagent kits (V3.1; 10X Genomics, PN-1000268). The libraries were sequenced on a NovaSeq X plus sequencer (Illumina) with 100 cycles.

#### Analysis of scRNA-seq data

RNA reads sequenced from the scRNA-seq library were aligned to GRCh38 genome using Cell Ranger software (v.9.0.0; 10X Genomics) to generate raw gene counts. Reads mapped to pre-mRNA were counted to include un-spliced nuclear transcripts. Cell barcodes were called using Cellranger-9.0.0 default parameters. Then, the obtained count matrices were corrected for ambient mRNA using CellBender (v.0.3.0) with half the default learning rate. Read_CellBender_h5_Mat() function was used to extract sparse matrix with corrected counts from CellBender h5 output file.

#### Slingshot Pseudotime Trajectory Analysis

Pseudotime trajectories were inferred using the Slingshot R package. The normalized Seurat object was converted into a SingleCellExperiment and embedded in UMAP space after batch correction and clustering. Slingshot was applied using Seurat-derived clusters, with the trajectory constrained to start from the OPC cluster (PDGFRA+) and extend toward differentiated oligodendrocyte states. Pseudotime values assigned to each cell were used for downstream analyses, including density plots and box plots.

#### Statistical analysis

The sample size for each experiments was determined on the basis of previous publications^27,93^. Statistical analyses were performed using R v.4.1.0 (R Foundation for Statistical Computing, Vienna, Austria) or GraphPad Prism 9 (Graphpad, San Diego, California) as indicated in the legends and Methods. Data distribution was assumed to be normal if not otherwise stated. *P* < 0.05 was considered statistically significant.

#### Data availability

The single-nucleus RNA-seq dataset generated in this study has been deposited in the NCBI Gene Expression Omnibus (GEO) under accession number GSE315101. The bulk RNA-seq dataset generated in this study has been deposited in GEO under accession number GSE315100.

### Supplementary Figures

**Supp Fig. 1.**
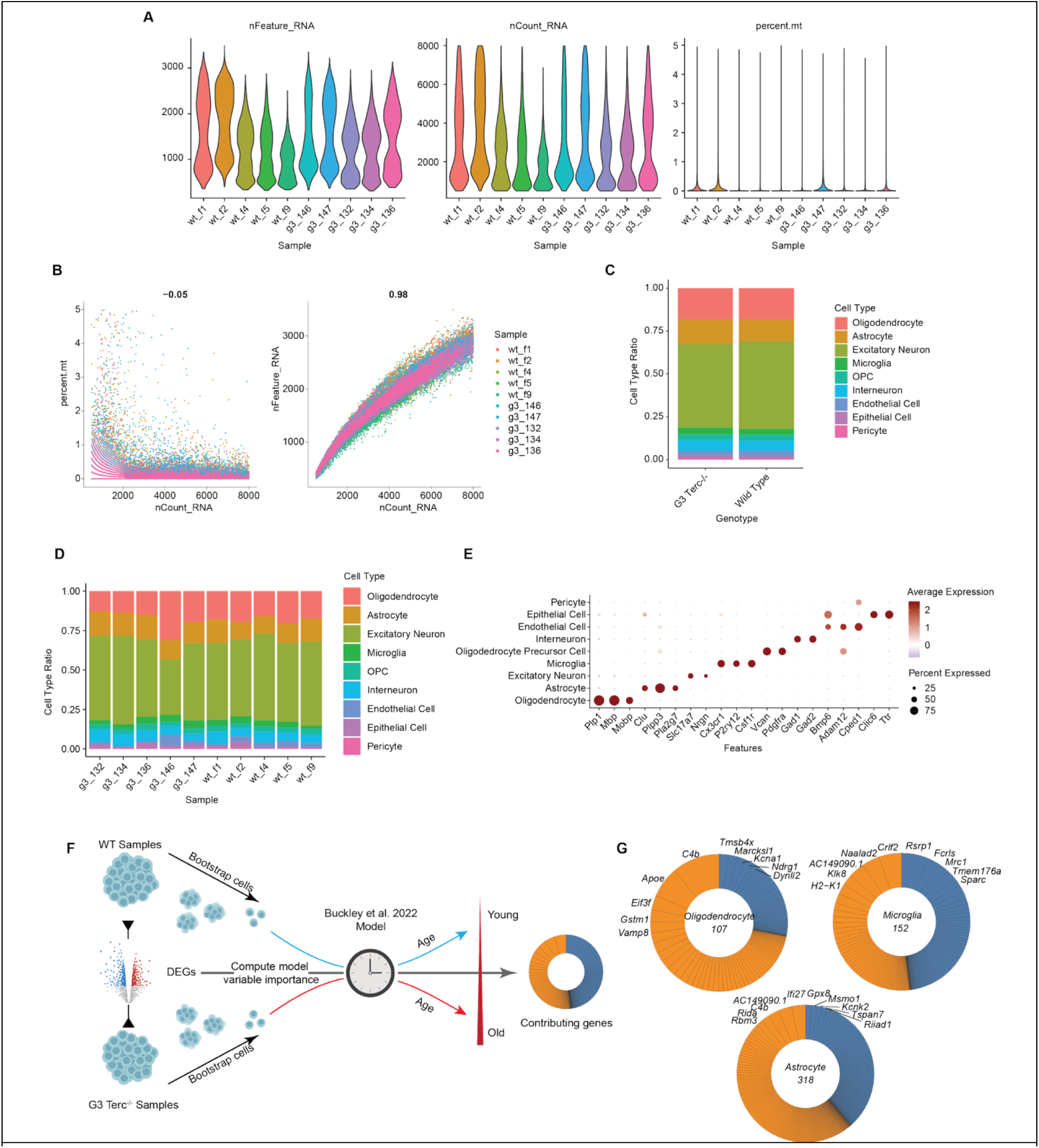
Quality control assessment of single-nuclei RNA-Seq in the G3 Terc^-/-^ cohort and the age-predicting model for glial cells. Related to Figure 2. **A.** Violin plots showing the number of unique features (left); the number of total RNA count (middle), and the percentage of mitochondrial genes (right) detected in each sample. **B.** Correlation between UMI counts and percentage of mitochondrial genes (left) or total gene counts (right) per nuclei for each individual sample. **C.** Proportion of each cell type within WT and G3 Terc^-/-^ genotypes. **D.** Proportion of each cell type within individual samples. **E.** Dot plot showing expression levels of canonical cell markers in each identified cell types. **F.** Schematic showing the application of aging-predicting model on G3 Terc-/- snRNA data. **G.** Donut plots showing the weight of genes contributing to the prediction of chronological age in oligodendrocytes, astrocytes, and microglia. The orange segments represent positive correlation to predicted age, and the blue segments represent negative correlation.

**Supp Fig. 2.**
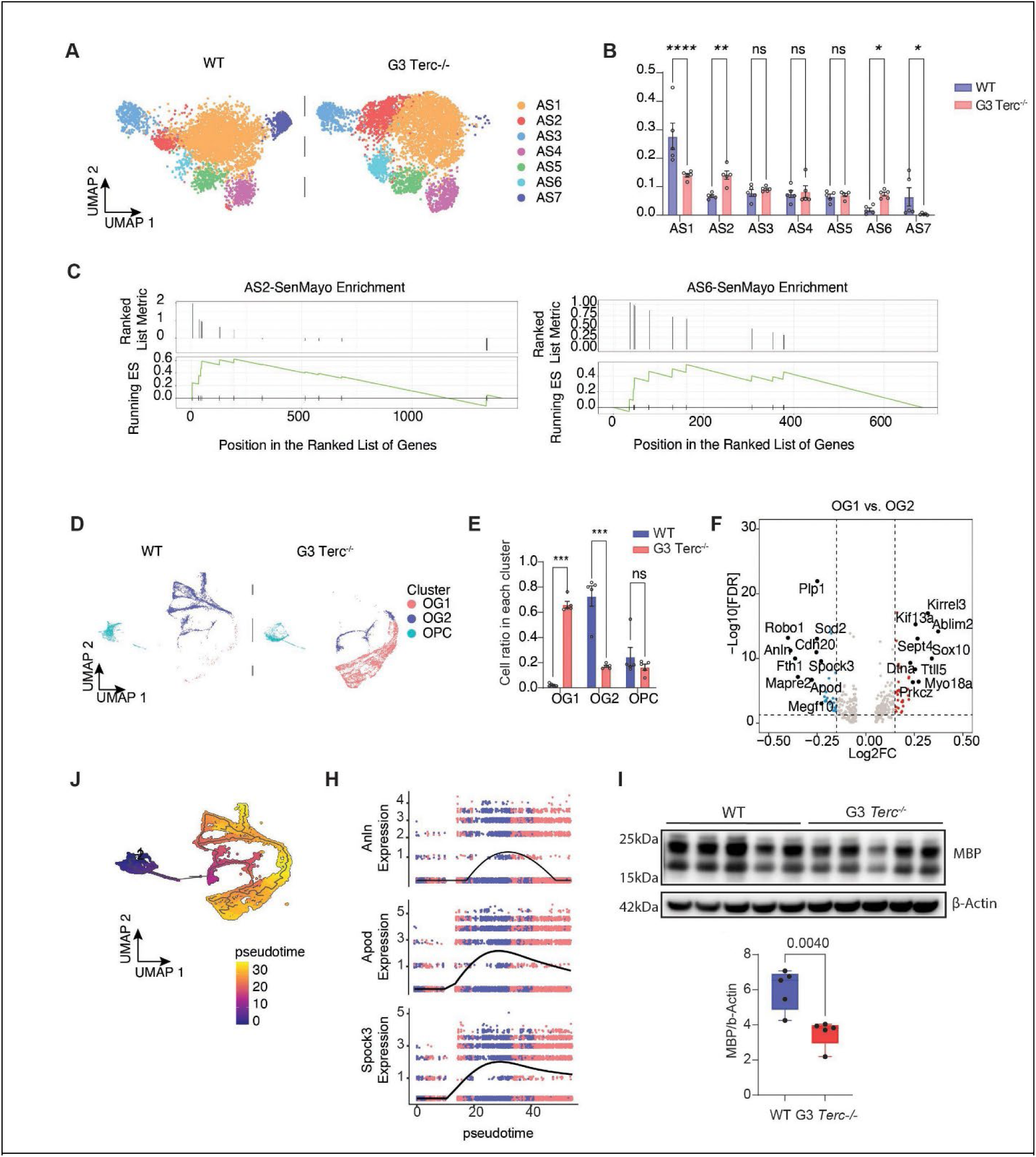
Telomere shortening induces senescent pathway in astrocytes and disrupts myelination. Related to Figure 2. **A.** UMAP plot showing a shift in astrocyte subclusters by G3 Terc^-/-^. **B.** Ratios of each astrocyte subcluster. Data are reported as mean ± s.e.m. and analyzed by two-way ANOVA. \*\*\*\**p*<0.0001; \*\**p*<0.005. ;\**p*<0.05. **C.** Running enrichment score and pre-ranked list showing a positive enrichment of the SenMayo gene set predicted by the upregulated DEGs in AS2 (left) and AS6 (right). **D.** UMAP plot showing a shift from OG2 (blue) to OG1 (red) caused by G3 Terc^-/-^. **E.** Ratios of each oligodendrocyte lineage cell cluster. Data are reported as mean ± s.e.m. and analyzed by two-way ANOVA. \*\*\**p*<0.0001 **F.** Volcano plot of DEGs comparing OG1 to OG2. Red and blue dots represent genes with a log2FC > 0.15 or < -0.15, respectively. All other genes are colored gray. The top differential genes are labeled. **G.** Pseudo-time trajectory depicting the development of oligodendrocyte from oligodendrocyte precursor cells. **H.** Expression dynamics of Anln, Apod, and Spock3 as a function of pseudo-time. **I.** Western blot of MBP in the cortex (upper panel), and **q**uantification of MBP signal normalized to β-actin (lower panel). Data were analyzed by two-tailed unpaired *t*-test.

**Supp Fig. 3.**
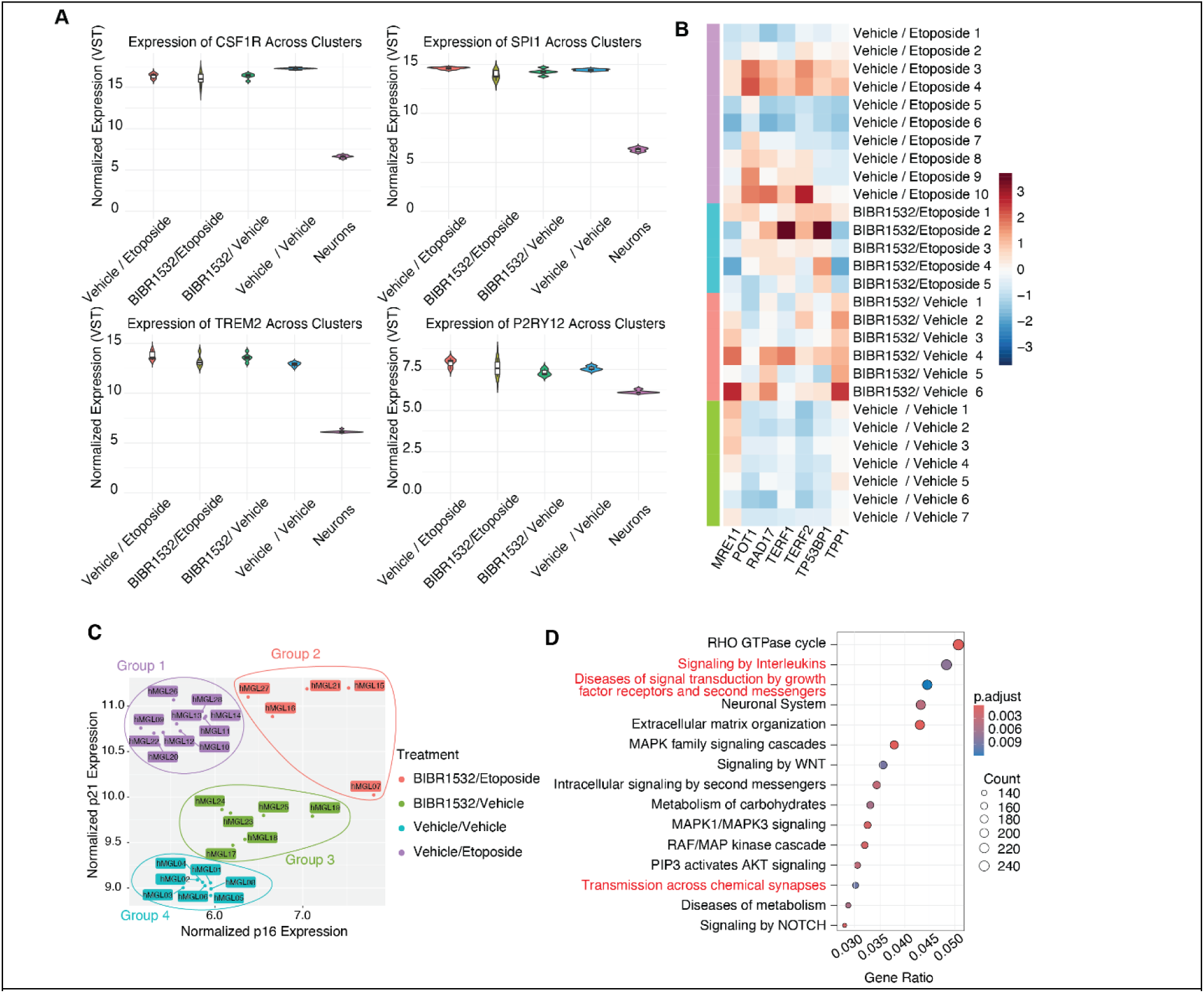
Quality control of hMGLs and BIBR1532/Etoposide treatment. Related to Figure 5. **A.** Violin plots showing the expression levels of the key microglia markers in hMGLs with different treatment conditions and neurons. **B.** Heatmap summary of the TIFs genes expressed in hMGLs with different treatment conditions and neurons. **C.** Scatter plot showing the expression levels of p16 and p21 in all RNA samples. Samples are clustered by k-means. **D.** Differentially regulated Reactome pathways predicted from senescent DEGs.

**Supp Fig. 4.**
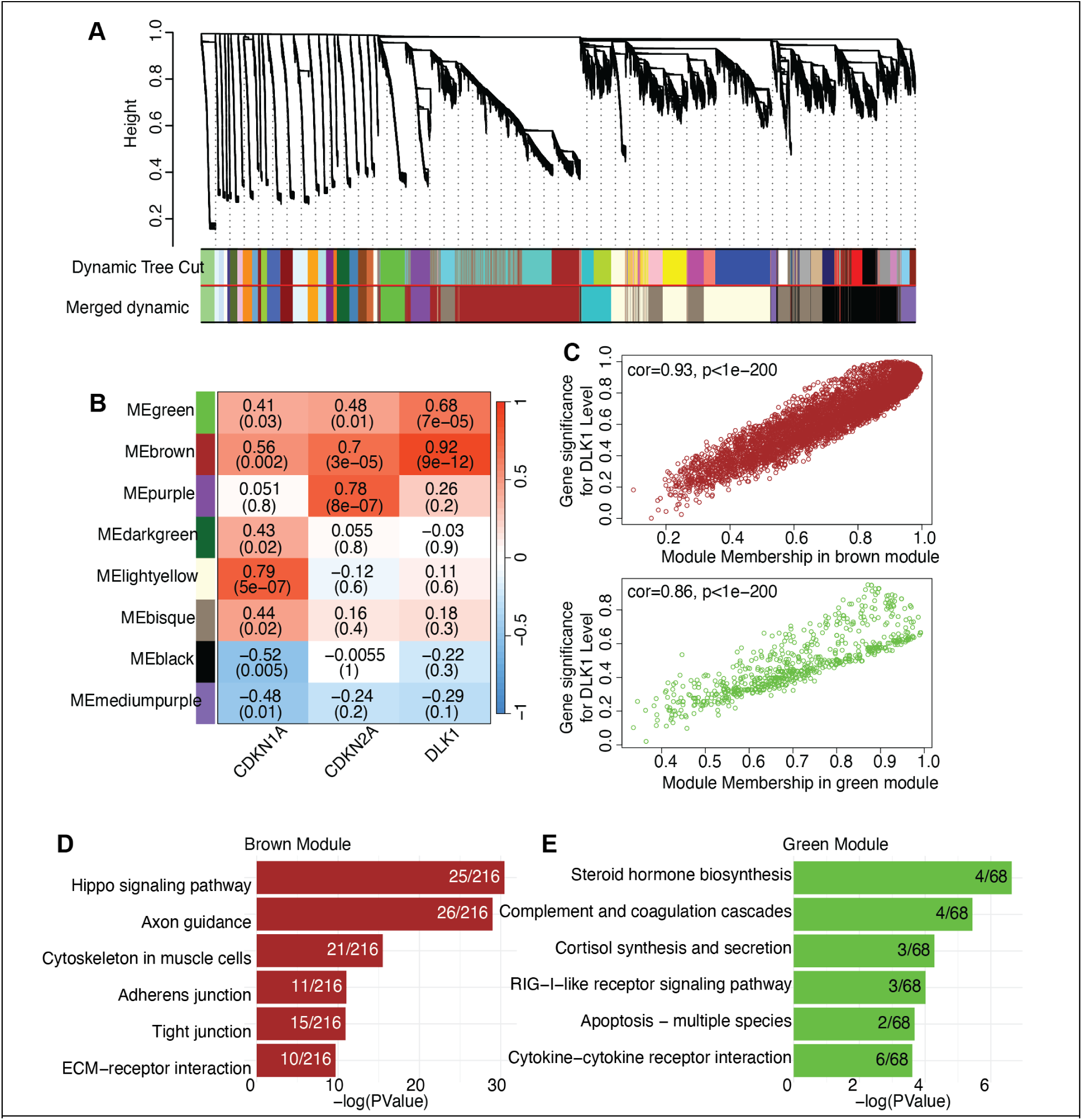
Gene networks associated with microglial senescence and DLK1 upregulation in hiMGLs. Related to Figure 5. **A.** Gene dendrogram showing all the gene modules identified in the iPSC-derived microglia transcriptome data. The merged dynamic was generated using a height threshold of 0.5. **B.** Heatmap showing the WGCNA modules significantly (p <= 0.05) correlated with p21, p16, or DLK1. **C.** Scatterplot of gene significance for DLK1 expression level versus module membership in the brown module (top) and the green module (bottom). **D–E.** Top six KEGG (Kyoto Encyclopedia of Genes and Genomes) pathways predicted from the top 500 genes (ranked by module membership) in the brown (D) and green module (E).

**Supp Fig. 5.**
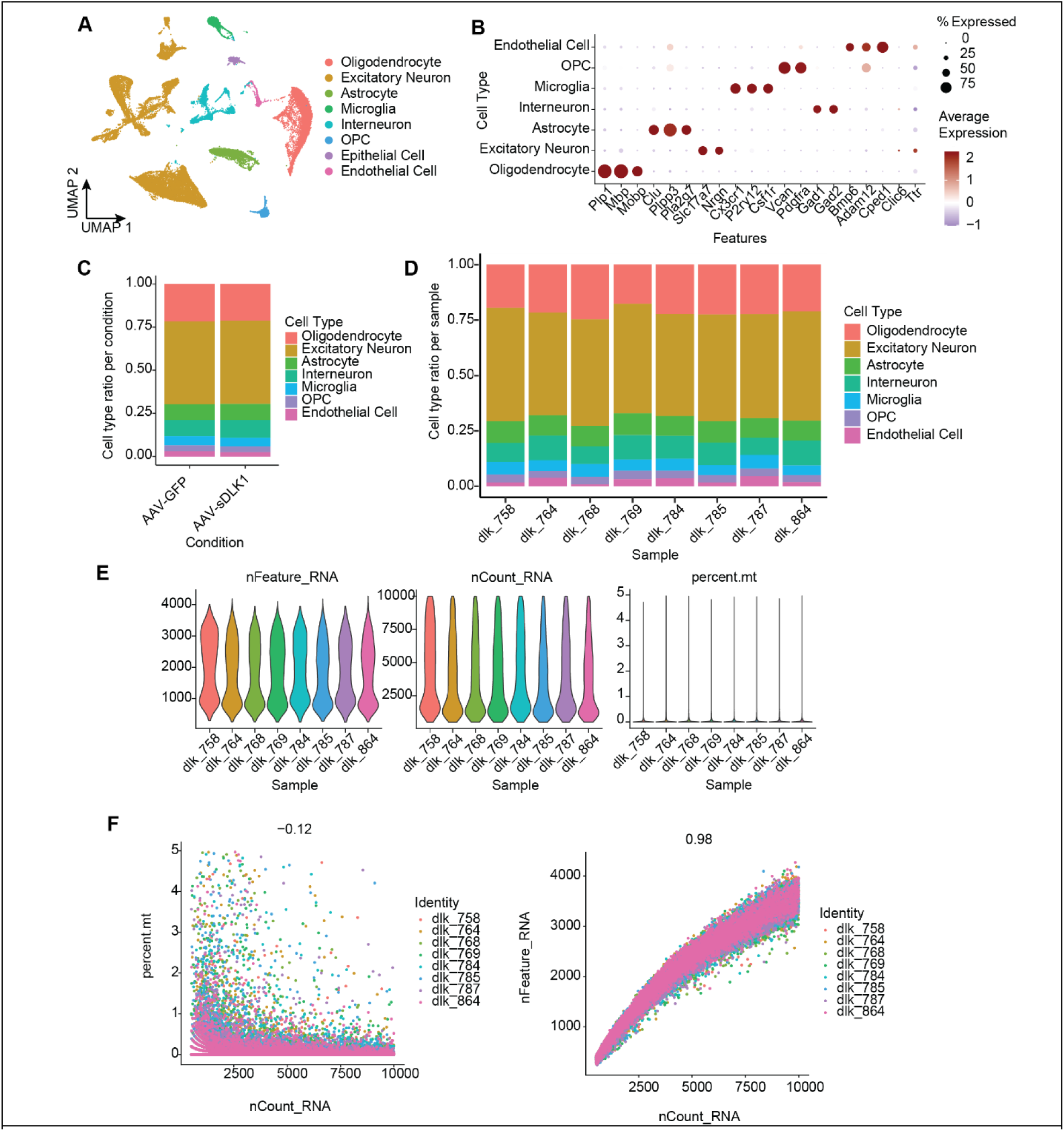
Quality control assessment of single-nuclei RNA-Seq in the AAV-sDLK1-T2A-GFP cohort. Related to Figure 6. **A.** UMAP plots showing eight major cell types identified in the mouse hippocampus. **B.** Dot plot showing expression levels of canonical cell markers in each identified cell type. **C.** Proportion of each cell type within animals injected with AAV-GFP and AAV-sDLK1-T2A-GFP. **D.** Proportion of each cell type within individual samples. **E.** Violin plots showing the number of unique features (left); the number of total RNA count (middle), and the percentage of mitochondrial genes (right) detected in each identified cell type. F. Correlation between UMI counts and percentage of mitochondrial genes (left) or total gene counts (right) per nuclei for each individual sample.

**Supp Fig. 6.**
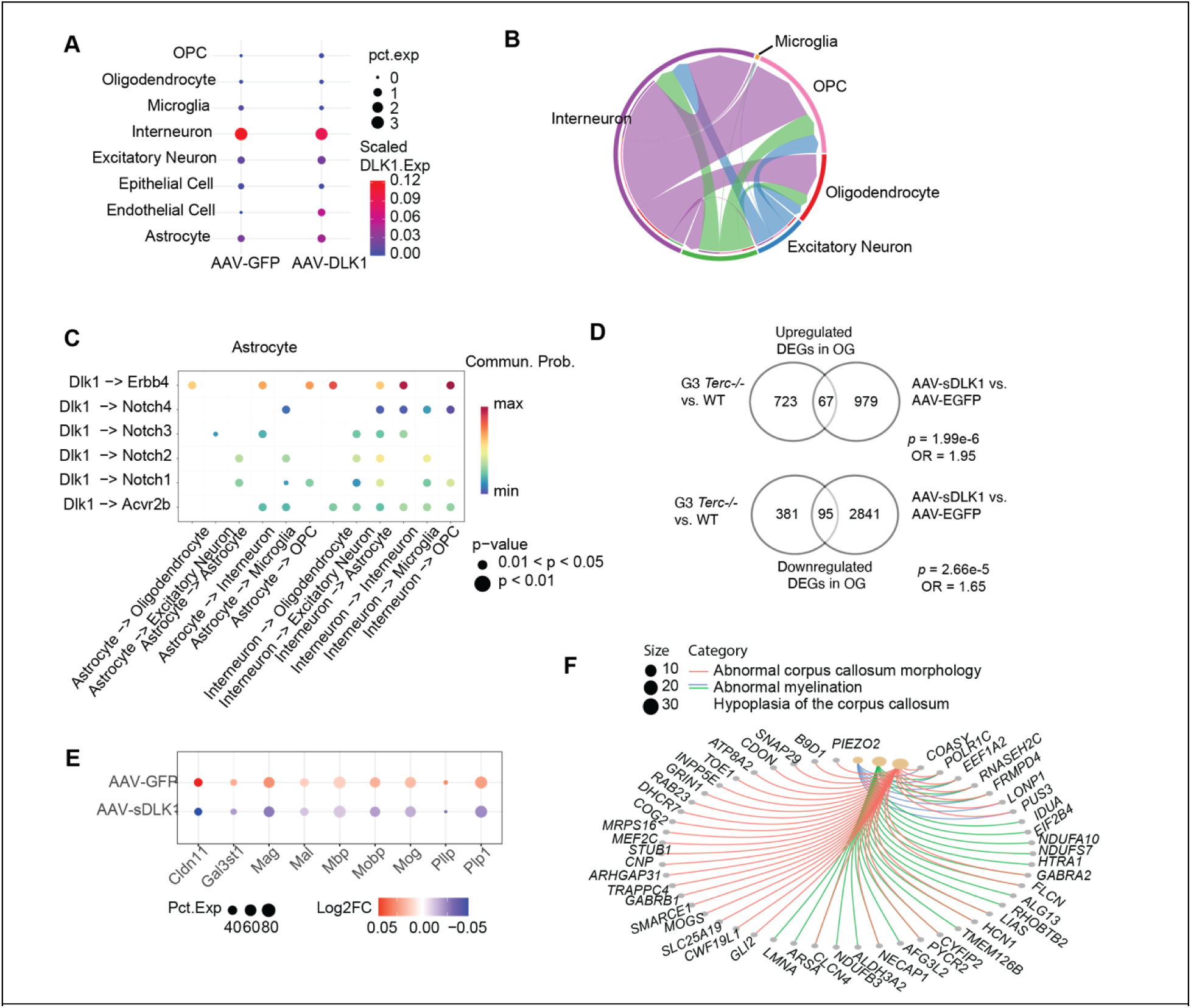
sDLK1 disrupts oligodendrocyte maturation and myelination. Related to Figure 6. **A.** Dot plot showing the expression level of DLK1 in different cell types in the hippocampus tissue. The size of each dot represents the percentage of cells with detected DLK1 mRNA. **B.** Chord diagram showing DLK signaling predicted by CellChat. The lengths of the segmented outer circle reflect the expression levels of ligand proteins in each cell type and of receptor proteins in the receiving cells, showing strong expression of DLK1 signaling originating from interneurons and astrocytes to oligodendrocytes and OPCs. **C.** Bubble plot showing the DLK1 interactions originating from interneurons and astrocytes to different receptors. **D.** Venn diagram showing the overlap of upregulated DEGs in G3 Terc-/- and AAV-sDLK1-T2A-GFP injected oligodendrocyte(top) and the overlap of downregulated DEGs in G3 Terc-/-and AAV-sDLK1-T2A-GFP injected oligodendrocyte (bottom). **E.** Dot plot showing the change of expression levels of myelination proteins in oligodendrocytes caused by the increase of sDLK1. **F.** Cnet plot showing the network of genes associated with myelination-related Gene Ontology terms and myelin-related diseases, based on enrichment analysis of the top 500 differentially expressed genes in oligodendrocytes from mice with AAV-sDLK1-T2A-GFP versus mice with AAV-GFP. Nodes represent genes or GO terms; edge colors represent the pathways each node is involved in.

**Supp Fig. 7.**
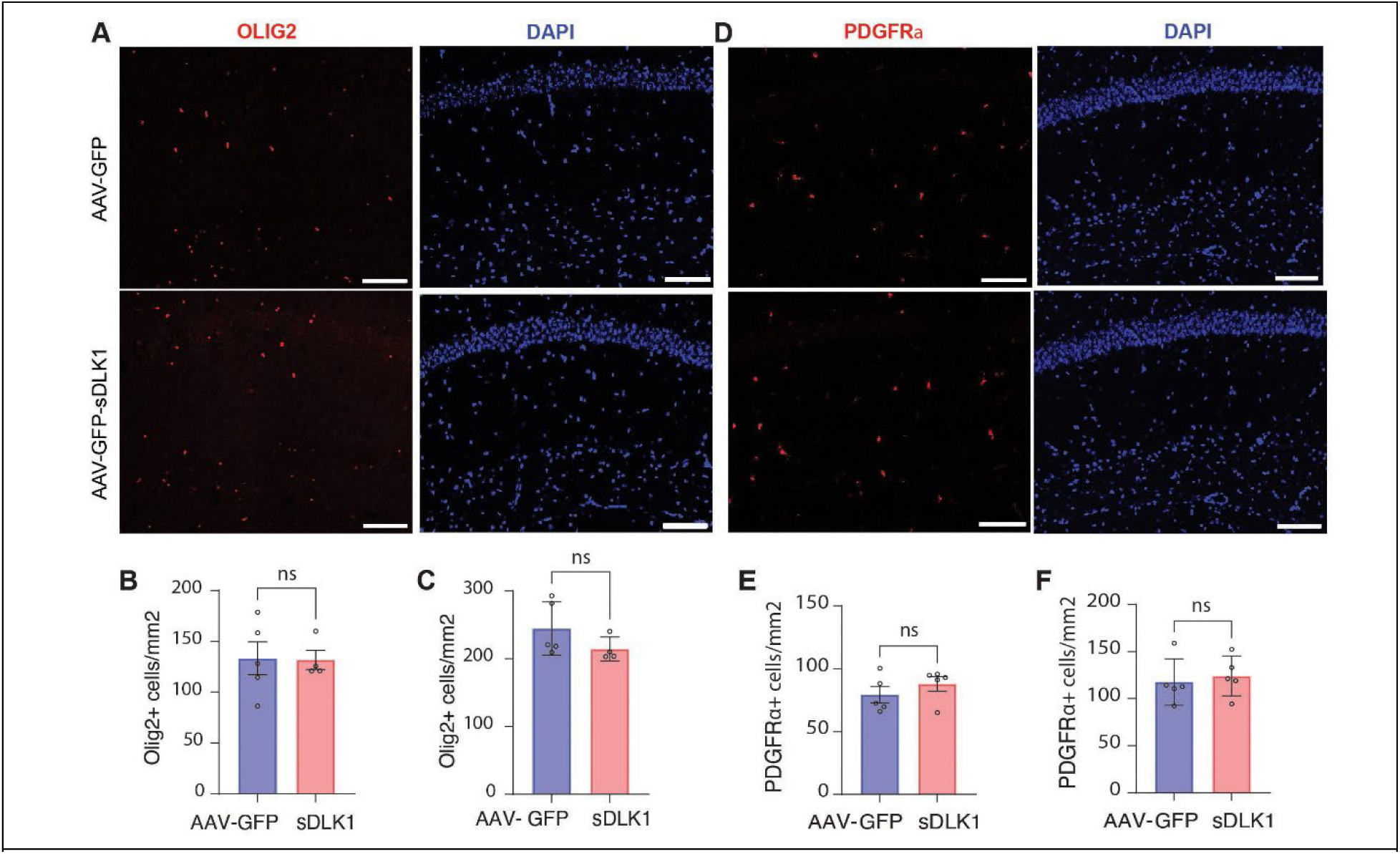
Elevated sDLK1 does not affect the number of OLs or OPC in the hippocampus. Related to Figure 6. **A.** Representative 20X images of Olig2 in the CA1 region of the hippocampus in C57BL/6 mice injected with either AAV-GFP or AAV-sDLK1-T2A-GFP. Scale bar represents 100 µm. **B.** Quantification of Olig2+ cell density in CA1. N = 5 mice injected with AAV-GFP and N = 4 mice injected with AAV-sDLK1-T2A-GFP. 2-3 hippocampal sections/mouse were imaged and analyzed. Data are reported as mean ± SEM. p = 0.9480. Data were analyzed by unpaired t-test. **C.** Quantification of Olig2+ cell density in CA3. N = 5 mice injected with AAV-GFP and N = 4 mice injected with AAV-sDLK1-T2A-GFP. 2-3 hippocampal sections/mouse were imaged and analyzed. Data are reported as mean ± SEM. p = 0.2008. Data were analyzed by unpaired t-test. **D.** Representative 20X images of PDGFRα in the CA1 region of the hippocampus in C57BL/6 mice injected with either AAV-GFP or AAV-sDLK1-T2A-GFP. Scale bar represents 100 µm. **E.** Quantification of PDGFRα+ cell density in CA1. N = 5 mice/treatment group and 2-3 hippocampal sections/mouse were imaged and analyzed. Data are reported as mean ± SEM. p = 0.3534. Data were analyzed by unpaired t-test. **F.** Quantification of PDGFRα+ cell density in CA3. N = 5 mice/treatment group and 2-3 hippocampal sections/mouse were imaged and analyzed. Data are reported as mean ± SEM. p = 0.6744. Data were analyzed by unpaired t-test.

**Supp Fig. 8.**
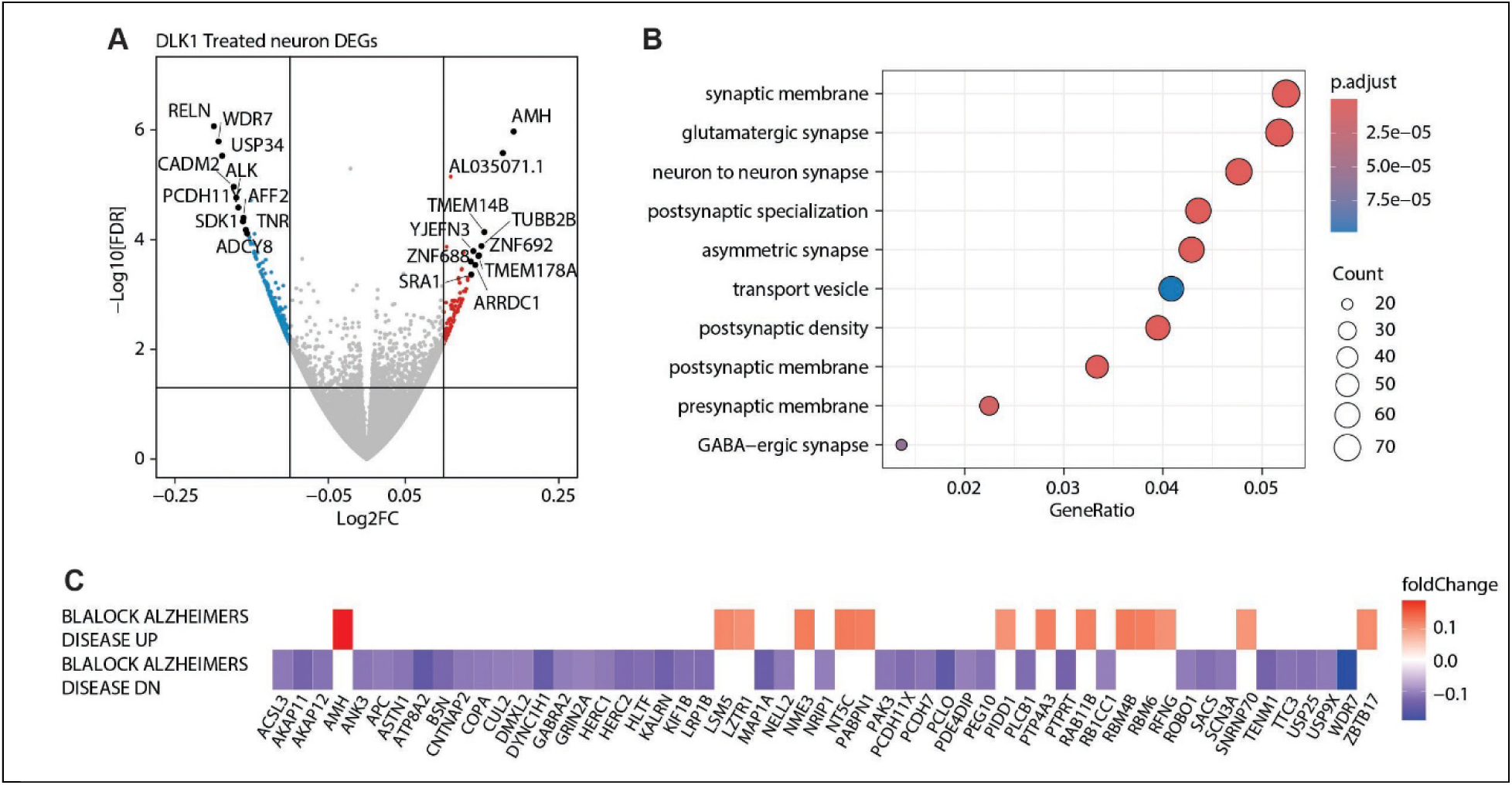
Soluble DLK1 alters the transcriptions of synaptic proteins in the iPSC-derived neurons. Related to Figure 8. **A.** Volcano plot of significant DEGs in iPSC-derived neurons treated with soluble DLK1. Red and grey dots represent genes with a log2FC > 0.1 or < -0.1, respectively. Top 10 differential genes are labeled. **B.** Dot plot of the top 10 GO Cellular Component ontologies inferred by the upregulated overlapping DEGs in sDLK1-treated neurons. **C.** Heatmap showing the fold-change of the genes upregulated (top) and downregulated (bottom) in the brains from patients with Alzheimer’s disease. All gene expressions are significantly changed (p-value <= 0.05) in the sDLK1-treated neurons.

